# Spatially resolved host-bacteria-fungi interactomes via spatial metatranscriptomics

**DOI:** 10.1101/2022.07.18.496977

**Authors:** Sami Saarenpää, Or Shalev, Haim Ashkenazy, Vanessa de Oliveira-Carlos, Derek Severi Lundberg, Detlef Weigel, Stefania Giacomello

**Affiliations:** SciLifeLab, Department of Gene Technology, KTH Royal Institute of Technology, Stockholm, Sweden; Max Planck Institute for Biology Tübingen, Tübingen, Germany; University of Tübingen, Tübingen, Germany; Zellbiologie und Epigenetik, Technische Universität Darmstadt, Darmstadt, Germany; Swedish University of Agricultural Sciences, Uppsala, Sweden

## Abstract

All multicellular organisms are closely associated with microbes, which have a major impact on the health of their host. The interactions of microbes among themselves and with the host take place at the microscale, forming complex networks and spatial patterns that are rarely well understood due to the lack of suitable analytical methods. The importance of high-resolution spatial molecular information has become widely appreciated with the recent advent of spatially resolved transcriptomics. Here, we present Spatial metaTranscriptomics (SmT), a sequencing-based approach that leverages 16S/18S/ITS/poly-d(T) multimodal arrays for simultaneous host transcriptome- and microbiome-wide characterization of tissues at 55-µm resolution. We showcase SmT in outdoor-grown *Arabidopsis thaliana* leaves as a model system, and found tissue-scale bacterial and fungal hotspots. By network analysis, we study inter- and intra-kingdom spatial interactions among microbes, as well as the host response to microbial hotspots. SmT is a powerful new strategy that will be pivotal to answering fundamental questions on host-microbiome interplay.

---

Advances in spatially resolved transcriptomics technologies have greatly improved the understanding of eukaryotic host gene expression mechanisms in animal and plant tissues (Eng et al., 2019; Giacomello et al., 2017; Rodriques et al., 2019; Ståhl et al., 2016). These technologies have been designed to capture targeted (Chen et al., 2015; Eng et al., 2019; Ke et al., 2013) or untargeted (Giacomello et al., 2017; Rodriques et al., 2019; Ståhl et al., 2016) RNA information based on imaging or sequencing of unique molecules, enabling the study of hundreds of genes or the whole transcriptome, respectively.

Spatial variation is also prominent in host-microbe interactions, and single-cell RNA-sequencing (scRNA-seq) of the host has been used to understand how this affects host cellular responses during infection (Tian et al., 2022). However, integrated spatially resolved analyses of microbial identity and the host response remain rare, and are typically focused on individual microbial taxa within a host (Sounart et al., 2022). Importantly, with existing technology it has not been possible to simultaneously resolve the spatial interactions between a host and the multitude of microbes colonizing it. This has considerably limited our understanding of host-microbe interactions at the tissue level.

Microbes often live in diverse communities surrounded by other microbes. Both cooperative and antagonistic interactions between microbes are known to be important for the functionality and health of ecosystems, plants, animals and humans (Durán et al., 2018; Fan and Pedersen, 2021; Logares et al., 2020). Moreover, the success of microbial colonization and infection depends strongly on the spatial structure of microbial interactions with other microbes and with multicellular species, and several pioneering studies have revealed clear and functionally significant spatial organization in host-associated microbial communities (Kim et al., 2020; Mark Welch et al., 2016, 2017). Much broader knowledge of the spatial organization of microbes within hosts, and the associated local host responses are therefore needed to fully understand the biology of the host-microbe-microbe interactome.

Fluorescence-based FISH techniques provided the first insights into microbial spatial organization in different environments (Dar et al., 2021) and in host tissues including: mouse gut (Dar et al., 2021), human plaque microfilms(Shi et al., 2020), and *Arabidopsis thaliana* roots (Cao et al., 2021). A limitation of these targeted methods is that they use a set of predesigned probes, each specific to a single microbial taxon. Current FISH-based technologies thus cannot provide comprehensive spatial descriptions of unknown microbiomes. Moreover, despite recent advances, these methods cannot yet achieve complete spatial resolution of the host’s expression patterns due to their limited capacity and overfitting to specific hosts (Xia et al., 2019).

Plants are colonized by a heterogeneous sets of microbes whose diversity is comparable to that of the human gut’s microbial population (Hacquard et al., 2015). Similar to gut microbes, plant colonizing microbes affect the host’s health and physiology in various ways, ranging from beneficial (Finkel et al., 2017) to harmful (Dean et al., 2012; Mansfield et al., 2012). Plant microbial communities are shaped in an environment-dependent manner by the intertwined forces of host-microbe and microbe-microbe interactions, which ultimately determine the fitness of the host and the associated microbes (Penczykowski et al., 2016).

Because of the limitations of current analytical methods, microbial interactions within plants are often deduced from complete tissues or whole plants, based on 16S rDNA abundance data (Agler et al., 2016; Durán et al., 2018; Shalev et al., 2022). This approach inevitably makes it impossible to resolve microscale differences in abundance. Hence, bulk RNA-seq can only be used to study average plant-microbe interactions in a tissue (Nobori et al., 2020; Vogel et al., 2016). Given the tremendous variation of unique RNA profiles found within tissues, demonstrated repeatedly by spatial transcriptomics and scRNA-seq analyses (Giacomello et al., 2017; Tabula Muris Consortium, 2020), it is very likely that important information has been obscured by the limited spatial resolution of the techniques used to study plant-microbe interactions.

Here, we present Spatial metaTranscriptomics (‘SmT’, **Figure 1**), an untargeted approach that allows simultaneous interrogation of bacterial and fungal communities, and the corresponding host transcriptional responses with a spatial resolution of 55 µm. By capturing the spatial distribution of bacterial and archaeal 16S rRNA sequences, together with fungal internal transcribed spacer (ITS) and 18S rRNA sequences and the host mRNAs, we link local changes in host gene expression to the size and composition of local microbial populations in *A. thaliana* leaves. We resolve the organization of microbial communities along tissue sections and demonstrate the presence of microbial hotspots at the leaf scale, and how these locally impact host responses.

**Figure 1:**
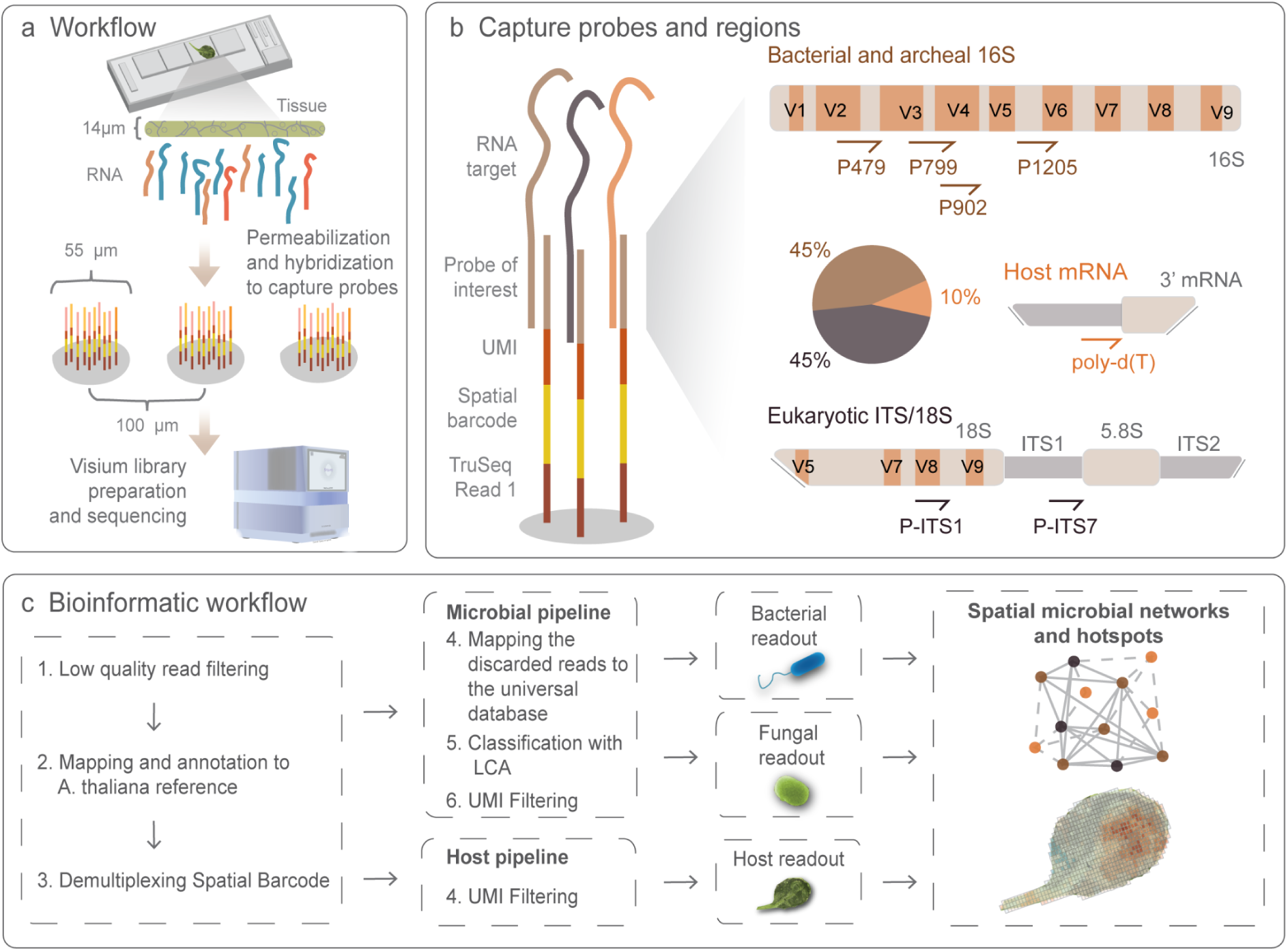
Overview of the method. **a.** Spatial metaTranscriptomics (SmT) uses capture arrays on glass slides. Each capture array contains 4,992 spots that are 55 µm in diameter and 100 µm from center to center. Cells are permeabilized to release RNA molecules that hybridize to the barcoded capture probes in the spots. The captured molecules are then processed into a sequencing library. **b.** Capture probes consist of a sequencing adapter, a spatial barcode, a unique molecular identifier (UMI), and a capture moiety. Poly-adenylated mRNAs are captured with poly-d(T) probes that comprise 10% of all the capture probes. Ribosomal RNAs from fungi are captured with P-ITS7 and P-ITS1 probes targeting the 18S rRNA and ITS regions, respectively. Ribosomal RNAs from bacteria and archaea are captured with P479, P799, P902, and P1205 probes targeting bacterial 16S rRNA. Bacterial and archaeal probes and fungal probes each comprise 45% of the capture probes. **c**. A bioinformatic workflow designed to assign the reads to host or microbial modalities. First, low-quality reads are filtered out, the remaining reads are mapped against the *A. thaliana* TAIR10 reference genome, and spatial barcodes are demultiplexed. Second, mapped *A. thaliana* reads are filtered based on their UMI and compiled to obtain a gene-count matrix. Third, the reads not mapping to *A. thaliana* are mapped to a universal database to remove those that are not clearly of microbial origin. The remaining microbial reads are classified with LCA based on their identity and UMIs, and unique taxa are counted to generate separate unique taxa-count matrices for fungi and for bacteria and archaea.

## Results

### Spatial detection of bacterial infection patterns and host response

To determine whether mRNA molecules could be captured from the host *A. thaliana* leaf sections while preserving the tissue’s morphology, we applied an optimized Spatial Transcriptomics (ST) protocol to leaves grown under laboratory conditions(Giacomello and Lundeberg, 2018). To this end, we permeabilized a 14-μm thick longitudinal leaf section on a glass surface uniformly coated with poly-d(T) capture probes. Following cDNA synthesis with fluorophores, we obtained a fluorescent cDNA footprint (**Figure 2a**) whose morphology matched that of the original leaf, demonstrating that spatial host gene expression patterns can be obtained from longitudinal leaf sections. Next, since bacterial communities are typically characterized based on 16S rDNA sequences, we hypothesized that capturing 16S rRNA molecules could provide information on the spatial distribution of bacteria in host tissues. To prove this concept, we analyzed leaves of soil-grown *A. thaliana* plants infiltrated with the model pathogen *Pseudomonas syringae* pv. *tomato* DC3000 (Pst DC3000), which was genetically labeled to enable its fluorescence imaging in whole leaves (**Figure 2b**). The array used in the analysis contained two degenerate probes (P799 and P902) to capture bacterial and archaeal (hereafter ‘bacterial’) diversity from 16S rRNA hypervariable regions, together with poly-d(T) probes to capture host mRNA, mixed in the following proportions: 50% poly-d(T), 25% P799, and 25% P902.

**Figure 2:**
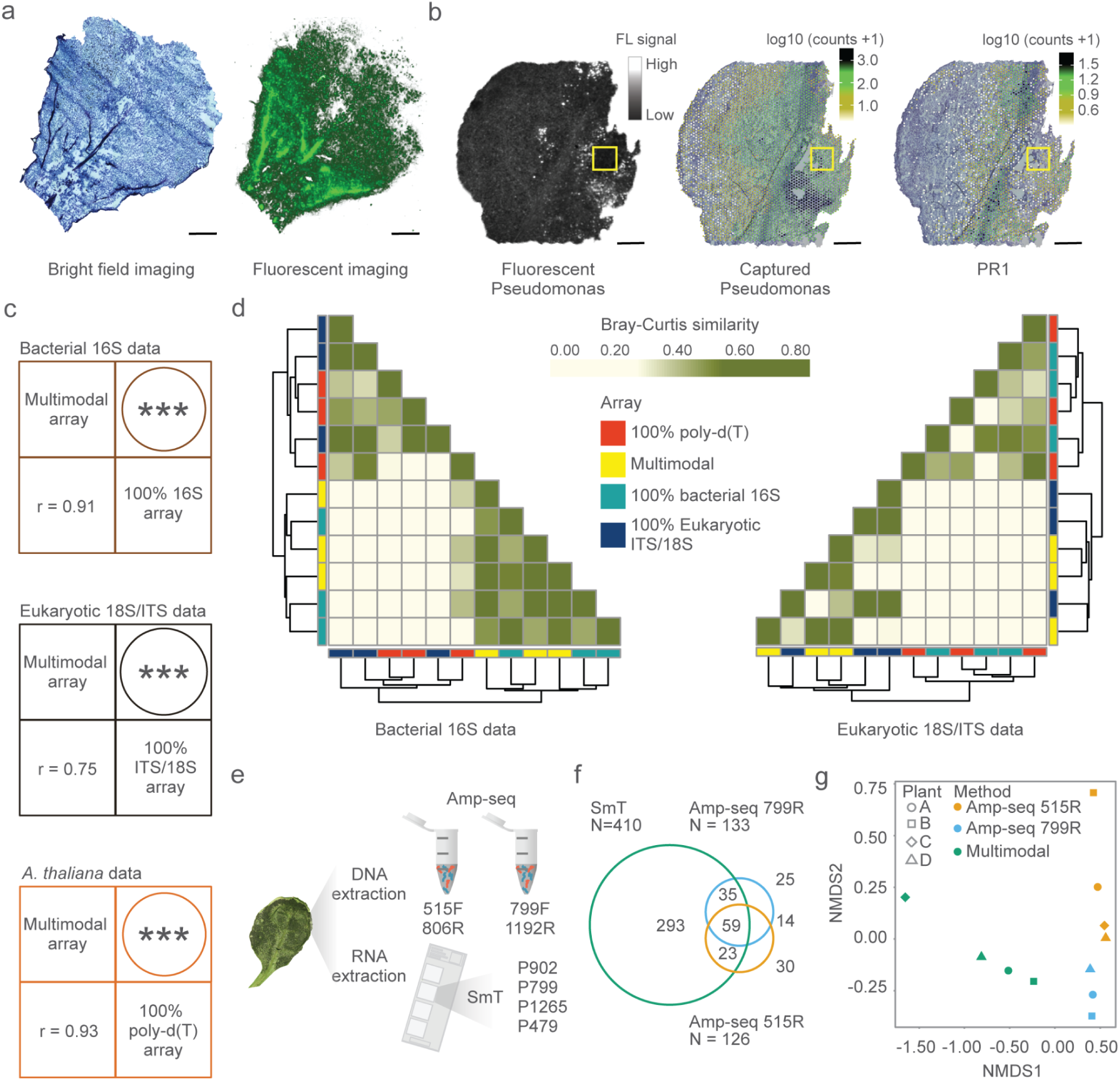
Spatial metaTranscriptomics resolves the microbial profile and host transcriptome at microscopic resolution. **a.** A Toluidine Blue-stained bright field image of a 14-µm thick longitudinal *Arabidopsis thaliana* leaf section (left) and the fluorescent cDNA footprint (right) of the same section from the tissue optimization experiment. **b.** Left: fluorescent image of an intact *A. thaliana* leaf syringe-infiltrated (yellow square) with mCherry-tagged Pst DC3000 bacteria. Middle: a 14-µm thick longitudinal section from the same leaf analyzed using a 50% poly-d(T) - 25% P799 - 25% P902 array, revealing the spatial capture of Pst DC3000 16S rRNA molecules. Right: spatial distribution of *PR1* gene expression in the same leaf section. **c.** Correlation of bacterial 16S rRNA, eukaryotic 18S rRNA/ITS, and *A. thaliana* molecules captured with a multimodal array containing 10% poly-d(T), 45% 16S, and 45% 18S rRNA/ITS probes and with 100% 16S rRNA, or 18S rRNA/ITS, and poly-d(T) arrays. **d.** Bray-Curtis similarity for bacterial and fungal taxa captured on different arrays, organized by hierarchical clustering. **e.** Experimental validation of SmT by amplicon sequencing. Numbers of bacterial and archaeal taxa detected using the two methods in a representative sample of four leaves from two plants are compared qualitatively using a Venn diagram (**f)** and quantitatively using non-metric multidimensional scaling (NMDS) (**g**).

We imaged intact infected leaves three days post-infiltration to record the fluorescent spatial pattern of the bacterial infiltration. and analyzed corresponding 14-µm-thick tissue sections with the array described above. We detected a uniform host and bacterial molecular capture throughout the tissue section (**Supplementary Figure S1**), indicating successful tissue permeabilization and RNA hybridization. We then compared the Pst DC3000 fluorescent signal to the distribution of 16S rRNA molecules (**Figure 2b**). Because the plants had been grown under non-sterile conditions, we expected to detect both Pst DC3000 and other bacteria. In total, we identified 512,779 unique bacterial molecules corresponding to 169 different bacterial taxa. The vast majority, 473,589 (92.4%), was identified as *Pseudomonas*, and these coincide with the areas where fluorescent signals were detected, indicating the ability to spatially capture the bacterial content. In addition, the density of *Pseudomonas* 16S rRNA molecules was highest at the infiltration site (indicated by the yellow squares in **Figure 2b**) and gradually declined towards distal regions, providing a more quantitative picture than the fluorescent imaging, which had missed the spatial component of the infection gradient.

We next investigated the expression patterns of host genes known to be involved in plant immune responses (**Figure 2b**). The spatial pattern of *PATHOGENESIS-RELATED GENE 1* (*PR1*), a marker of the plant immune response (Bretz et al., 2003), closely matched the distribution of *Pseudomonas*. Taken together, these results show that we are able to simultaneously capture bacterial taxonomic information and host transcripts.

### A multimodal array for simultaneous detection of microbial and host spatial information

Having demonstrated that bacterial information can be specifically captured together with information on host gene expression, we aimed to add a third modality to our arrays, capturing information from eukaryotic microbes, specifically fungi. We designed 18S rRNA/ITS probes specific for fungi and tested their performance in both separate arrays and a multimodal array. For this purpose, we created arrays with 100% poly-d(T) probes, 100% 16S rRNA probes, and 100% 18S rRNA/ITS probes, as well as a multimodal array containing all three probe types (10% poly-d(T), 45% 16S rRNA, and 45%18S rRNA/ITS). We dissected three leaves of outdoor-grown *Arabidopsis* plants into four 14-μm thick longitudinal sections, and analyzed consecutive sections from each leaf using the four array types. The multimodal and unimodal arrays greatly enriched the proportion of captured reads for the corresponding taxa when compared to the unspecific poly-d(T) probes (**Supplementary Figure S2**). Specifically, at the genus level, the multimodal array enriched bacterial and fungal unique molecules up to ∼19 and ∼31 fold, respectively. At the super kingdom level, the 100% 16S rRNA array enriched bacterial unique molecules up to ∼47-fold when compared to the 100% poly-d(T) array, and the 100% 18S rRNA/ITS array enriched fungal unique molecules up to ∼233-fold. As expected, the multimodal array enriched microbial signals to a lesser degree than the 100% 16S rRNA and 100% 18S rRNA/ITS arrays, given the lower concentration of microbe-specific probes in the multimodal arrays (**Supplementary Figure S2**).

Importantly, the bacterial information captured using the multimodal arrays was almost identical to that captured from consecutive tissue sections using 100% 16S rRNA arrays (both qualitatively and quantitatively). The multimodal array captured up to 962 bacterial taxa and 179 fungal taxa at the genus level (**Supplementary Table S1**), and recapitulated the profile of 100% 16S rRNA arrays independently if full bacterial components (r=0.90-0.92, P<0.001), top 500 bacterial taxa (r=0.91-0.93, P<0.001) or top 20 bacterial taxa (r=0.96-0.99, P<0.001) were considered (**Figure 2c** and **Supplementary Figure S3-S5**). Similarly, the multimodal array recapitulated the profile of 100% 18S rRNA/ITS arrays if full bacterial components (r=0.73-0.77, P<0.001) were considered, while the correlations obtained for the top 500 and 20 fungal taxa were 0.68 and 0.85 (P<0.001), respectively (**Figure 2c**, **Supplementary Figure S6-S8**).

Bray-Curtis similarity showed that the bacterial profile obtained using the bacterial 16S rRNA array was most similar to that of the multimodal array, while the fungal profile obtained with the multimodal array clustered with that for the eukaryotic 18S rRNA/ITS array (**Figure 2d**). Conversely, the bacterial profile obtained with the eukaryotic 18S rRNA/ITS array and the poly-d(T) array differed markedly from that obtained with the bacterial 16S rRNA array, and the fungal profile obtained with the bacterial 16S array and the poly-d(T) array differed markedly from that obtained with the 18S rRNA/ITS array. The multimodal array thus achieved high similarity to the 100% arrays for bacteria and fungi, but the same was not true when a kingdom-specific array was used to analyze a kingdom other than that for which it was designed. We confirmed this result by calculating the Shannon diversity index across leaves, revealing that the multimodal and 100% 16S rRNA arrays captured similar levels of diversity (H’=3.62-4.01 and H’=3.81-4.04, respectively), different from the 100% 18S rRNA/ITS and 100% poly-d(T) arrays (H’=2.76 and H’=3.70, respectively, **Supplementary Figure S9**). Overall, the bacterial profile captured by the 16S rRNA array and the fungal profile captured by the 18S rRNA/ITS array could only be recapitulated by the multimodal array and not by any of the unspecific probes (**Figure 2d**). These results imply that the multimodal array quantitatively enriched microbial counts and accurately profiled microbial populations within tissue sections, unlike the unspecific poly-d(T) probes (**Figure 2d**, **Supplementary Figure S2-S9**, **Supplementary Table S1**).

Finally, we confirmed that the multimodal array correctly captured the full transcriptomic profile of the host as well (**Figure 2c and Supplementary Figure S10**) by comparing the *A. thaliana* gene expression pattern captured with the multimodal array to that obtained with the 100% poly-d(T) array. The multimodal array captured 15,709 *Arabidopsis* genes on average and its correlation with the 100% poly-d(T) array was high (r=0.92-0.93, P<0.001). Overall, these results show that multimodal arrays enable accurate simultaneous capture of the host transcriptome, the bacterial profile, and the fungal profile.

### Validation of the multimodal array with amplicon sequencing

To improve the taxonomic classification of bacteria using the multimodal array, we introduced two additional 16S rRNA probes, P479 and P1265 (**Supplementary Figure S11**, **Figure 1**), and compared the results with those from 16S rDNA amplicon sequencing (‘amp-seq’) – current gold standard for bacterial profiling. Amp-seq involves PCR amplification of crude DNA extracts using a primer pair. Conversely, our multimodal array captures RNA fragments that are targeted by individual probes. We sampled three leaves from field-grown *A. thaliana* plants, and simultaneously extracted their RNA and DNA. We then analyzed the crude RNA extracts with the multimodal array containing the additional P479 and P1265 probes, and used the extracted DNA for amp-seq of two 16S loci with V3-V4 (primers 515F + 806R) and V4-V6 (primers 799F + 1192R) (**Figure 2e**).

We first qualitatively compared the bacterial profiles obtained using the multimodal array to those obtained with the two single pairs of 16S rDNA amp-seq primers by analyzing the presence or absence of every genus found by at least one of the three processes (**Figure 2f**, **Supplementary Figure S12**). SmT detected more than three times the total number of taxa detected by the two amp-seq primer pairs (**Figure 2f**), including ∼71% of the taxa detected by the amp-seq V4-V6 primers and ∼65% of the taxa detected by the amp-seq V3-V4 primers. The two amp-seq primer pairs overlapped in 55% of detected taxa.

We obtained similar results for the other two biological replicates (**Supplementary Figure S12)**, and a similar trend in quantitative analyses, comparing the relative abundances (**Figure 2g**). Thus, the microbial profiles retrieved by SmT are comparable to those obtained using the standard 16S rDNA amp-seq method.

In summary, these results confirm that our multimodal array accurately profiles bacteria in *A. thaliana* leaves and captures a more diverse taxonomic range than standard amplicon sequencing.

### Microbial hotspots within the host govern microbial interactions

The spatial distribution of the members of natural microbial communities within host leaves has been largely unknown. We therefore used SmT to investigate the microbial profiles of different leaf sections in wild *A. thaliana* leaves. The microbial profiles of the different sections were similar, reflecting the similarity of the environments in which the source plants were grown (Methods) and the reproducibility of our method **(Figure 3a)**. Considering only taxa with relative abundances above 1%, we identified 29 bacterial taxa and 23 fungal taxa at the genus level (Supplementary **Table S2**, **S3**). The relative abundances of different microbes did not vary greatly across sections, leaves, or whole plants. Analysis of the overall spatial distributions of bacterial and fungal genera **(Figure 3b and Supplementary Figure S13)** revealed that microbes were present across almost the entire leaf surface: unique bacterial molecules were detected in 99.9% of sampling spots at an average density of ∼277 molecules per million reads, while unique fungal molecules were detected in 97.5% of sampling spots at an average density of ∼261 molecules per million reads (**Supplementary Table S4**).

**Figure 3:**
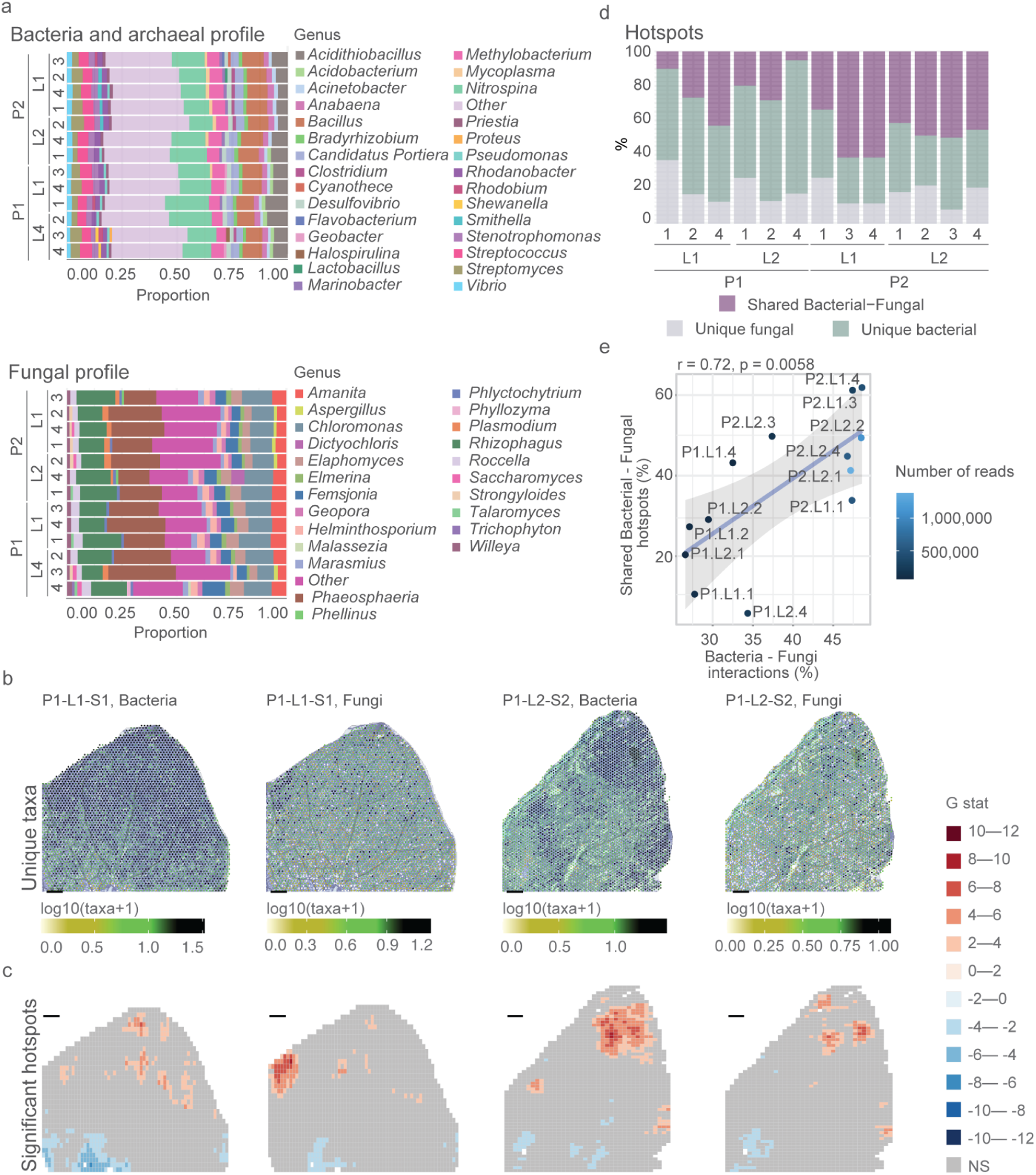
Microbial interactions are driven by spatial organization. **(a)** Bacterial and fungal profiles for each of the sections of four leaves (′L′) from two plants (′P′). ‘Other’ denotes binned bacterial and archaeal genera and fungal genera having <=1 % abundance. **(b)** Numbers of unique microbial taxa per capture spot and **(c)** significant hot- and cold-spots for bacteria and fungi in a representative leaf section. **(d)** Percentages of shared and unique hotspots among bacteria and fungi across 12 different leaf sections. Of note is the variance in shared regions across the sections. The sections were taken from 2 leaves from 2 different plants (4 leaves in total). ‘P’ - plant. ‘L’- leaf. **(e)** The proportion of interkingdom (bacteria-fungi) interactions as a function of the proportion of shared interkingdom hotspots.

We next analyzed the geography of microbial colonization. Although we detected both bacteria and fungi across the entire leaf surface, they were concentrated in hotspots rather than being homogeneously distributed (**Figure 3c; Supplementary Figure S14**). Some leaf regions were highly colonized with microbes, while others were uncolonized or colonized at very low levels. Further investigation revealed that some hot spots were shared between bacteria and fungi (**Figure 3d**, **Supplementary Figure S14**). The relative abundance of shared and unique hotspots varied widely across the 13 leaf sections (**Figure 3d**). Since microbial interactions are constrained by physical proximity (Esser et al., 2015), we hypothesized that the relative abundance of shared and unique hotspots control the proportion of inter- and intra-kingdom interactions. To test this, we computed the interaction network of the 50 most abundant bacterial and fungal taxa using an algorithm that accounts for the spatial structure of our data (Methods). We exemplify this approach by focusing on a subnetwork of 14 taxa (12 Bacterial and 2 Fungal) which are strongly associated (average pairwise Spearman’s rank correlation coefficient [SRCC]≥0.35) in all tested leaf sections (**Supplementary Figure S15**). We then tested the association between the relative abundance of shared hotspots and the magnitude of interkingdom (bacteria-fungi) interactions across the leaf sections, revealing a positive correlation between the two features (SRCC=0.72, P=0.0058; **Figure 3e**). This implies that microbial interactions are confounded by their spatial organization, and specifically by their presence in shared hotspots. We found similar association for the magnitude of bacteria-bacteria interactions and the fraction of bacterial-unique hotspots (SRCC=0.72, P=0.059; **Supplementary Figure S16**), but lower for fungi-fungi interactions and the fraction of fungal-unique hotspots (SRCC=0.47, P=0.1; **Supplementary Figure S17**).

Together, these results demonstrate a considerable spatial organization of microbes within the leaf.

### Microbial hotspots are associated with the local spatial gene expression of the host

Because microbe-microbe interactions are confounded by spatial relatedness, we hypothesized that microbial organization might also confound host-microbe interactions. We therefore investigated the effects of microbial hotspots on the host transcriptome by reducing the host expression patterns into 5 gene clusters using UMAP (**Figure 4a**, McInnes et al., 2018). As expected, the clustered spots reflected the leaf’s tissue structure, in which different cell types are distributed fairly evenly with the exception of vascular tissue. We identified mesophyll and vascular tissue-related gene clusters by considering gene annotations and through visual analysis (**Figure 4b**; **Supplementary Figure S18**). For example, *CHLOROPHYLL A/B BINDING PROTEIN 3* (*CAB3*), a common marker gene for mesophyll cells (Berkowitz et al., 2021), was upregulated in cluster 2 (avg. log_2_FC=0.33, **Supplementary Figure S19**). In addition, *GLUTATHIONE S-TRANSFERASE PHI 9* (*GSTF9*) was upregulated in cluster 3, which visually corresponded to leaf vasculature (**Supplementary Figure S19**) in agreement with data retrieved from the Single Cell Leaf Atlas (Kim et al., 2021).

**Figure 4:**
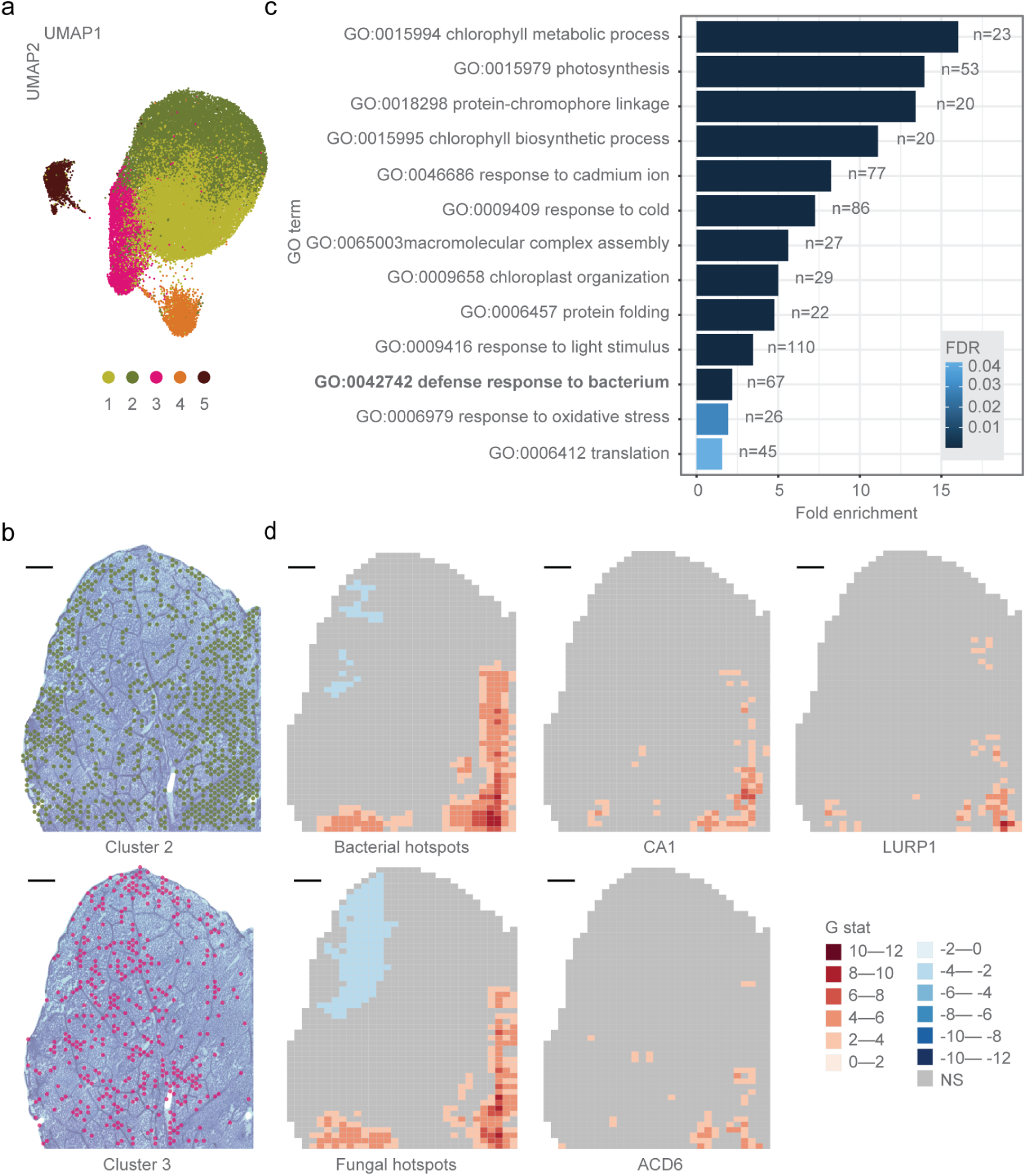
Host response is associated with microbial colonization pattern. **a**. UMAP clustering of host gene expression. **b**. Projection of UMAP clusters 2 and 3 onto the corresponding leaf section. **c**. Overrepresented GO terms for microbial-associated genes (n = number of genes labeled with the indicated GO term). **d**. Spatial distributions of significant bacterial and fungal hotspots together with hotspots for the expression of the defense-related genes *CA1* (AT3G01500), *LURP1* (AT2G14560), and *ACD6* (AT4G14400).

These results show that our system accurately resolves spatial host expression profiles in leaves. However, gene annotation analysis revealed no strong association between any of the five clusters and microbial colonization, and there was no obvious visual overlap between the clusters and the microbial hotspots (**Supplementary Figure S14**, **S18**).

To further investigate the host response to microbial hotspots, we first tested what fraction of expression hotspots overlapped with the microbial hotspots. We found that it highly varies across leaf sections, ranging from 16.7% to 75% shared expression-microbial hotspots (**Supplementary Figure S20**). Next, we performed a machine learning-based analysis (‘Boruta’) to associate the host’s spatial gene expression pattern with bacterial and fungal abundance (Methods). This revealed 1,323 and 954 host genes that were significantly associated with bacteria and fungi, respectively (**Supplementary Table S5**). To generalize our results, we focused on genes associated with microbial abundance in at least two sections of the same leaf (**Supplementary Figure S21**). This conservative approach reduced the number of genes associated with bacteria and fungi to 645 and 442, respectively. The vast majority of these (63% out of 667 genes in total) was associated with both kingdoms, indicating involvement in a general microbial response by the host, rather than a kingdom-specific one (**Supplementary Figure S22**). A Gene Ontology (GO) analysis revealed enrichment of biological process terms associated with plant immune responses, including GO:0042742–‘defense response to bacterium’ and GO:0006979–‘response to oxidative stress’ (**Figure 4c, Supplementary Figure S23, and Supplementary Table S6**). In total, 73 (11%) of the associated genes had GO terms associated with defense responses to bacteria and/or fungi (**Supplementary Table S7**). The spatial correlation between gene expression and microbial abundance is well illustrated by the expression patterns of three genes, *ACD6*, *CA1* and *LURP1* (**Figure 4d**, **Supplementary Figure S24**). All three genes are related to the basal plant immunity– *ACD6* is a broad-spectrum disease resistance gene activated by diverse microbes (Lu et al., 2003), the *CA1* gene product binds the immune-related hormone salicylic acid (Poque et al., 2018) and regulates stomatal opening during pathogen invasions (Zhou et al., 2020), and *LURP1* is required for resistance to the pathogen *Hyaloperonospora parasitica* (Knoth and Eulgem, 2008). Overall, these results reveal a connection between the spatial organization of microbes within the leaf and the host expression signature.

## Discussion

We present Spatial metaTranscriptomics (SmT), a multimodal untargeted sequencing method to investigate host-microbe-microbe interactions in tissue sections at a resolution of 55 µm. Numerous spatially resolved transcriptomics methods have been introduced so far (Moses and Pachter, 2022) based on either targeted (Eng et al., 2019; Merritt et al., 2020) or untargeted (Rodriques et al., 2019; Ståhl et al., 2016; Vickovic et al., 2019) capture of the transcriptional information and characterized by different spatial resolution ranging from subcellular (Chen et al., 2022, 2020; Xia et al., 2022) to multiple cells (Liu et al., 2020; Ståhl et al., 2016). These methods have been applied to a wide range of tissues from humans (Asp et al., 2019; Berglund et al., 2020; Hildebrandt et al., 2021) to plants (Duncan et al., 2016; Giacomello, 2021; Giacomello et al., 2017; Xia et al., 2022). Recently, methods have been developed that are capable of detecting multiple modalities such as protein and transcriptional information(Ben-Chetrit et al., 2022; Liu et al., 2020, 2022; Merritt et al., 2020; Wang et al., 2018) or chromatin accessibility and transcriptional information (Deng et al., 2022). SmT extends the reach of multimodal spatial measurement methods by capturing information from coexisting organisms.

SmT captures fungal, bacterial, and host signals from a tissue section while preserving their spatial structure and thus enabling integrated network analysis of gene expression by the host and its microbiota - prokaryotic and eukaryotic alike. Recent advances in smFISH techniques for microbiome analysis support the spatially resolved capture of over 1,000 bacterial taxa at the single cell level (Shi et al., 2020) or the detection of bacterial metabolic activities (Dar et al., 2021). However, smFISH is a laborious technique and requires the design of highly sensitive probes to capture a sample’s full bacterial diversity. Extending it to simultaneous detection of host whole-transcriptome information and potentially another microbial kingdom will likely be very challenging. SmT provides a straightforward approach, by sequencing the 16S rRNA and 18S rRNA/ITS variable regions together with polyadenylated transcripts. Our validation of SmT with amplicon sequencing, the gold standard method for bacterial profiling, revealed that SmT can more sensitively capture bacterial diversity than amplicon sequencing. Nevertheless, like any other emerging technologies, SmT presents limitations. By capturing information in spots of 55 μm in diameter, SmT does not achieve single-cell resolution. At least for the host, computational spot deconvolution methods could be applied to directly link cell type composition to a set of microbes.

We showcased SmT on *A. thaliana* leaves, which are an important model system for phyllosphere microbiology. We found microbial hotspots within plant leaves, reminiscent of microbial microniches in the human mouth <u>(Welch et al., 2020)</u>. An important question for future research will be whether there are physical constraints that favor a specific spatial organization of microbes within the leaf. We hypothesize that the invasion point at which the epiphytes entered the leaf is one factor governing the location of hotspots (Melotto et al., 2006), while the boundaries of hotspots may be set by the host response or simple ecological factors such as a local lack of nutrients in specific microenvironments (Geier et al., 2020). An ecologically important aspect of microbial hotspots is that interactions are strongest between microbes in close physical proximity (Esser et al., 2015; Tecon et al., 2018). New knowledge of inter-kingdom microbial interactions will be particularly valuable, given that inter-kingdom interactions can be associated with plant health (Agler et al., 2016; Durán et al., 2018).

As for microbial interactions, studies of plant responses to microbial colonization have mainly been limited to analyses of homogenized whole tissues (Finkel et al., 2020; Nobori et al., 2020; Vogel et al., 2016). SmT now allows us to link microbial abundance at the micrometer level to host transcriptional responses. We found a high degree of overlap between the sets of genes associated with bacteria and fungi, implying a general response of leaf cells to microbes, although this generality could be confounded by the extensive colocalization of bacteria and fungi in the sampled leaves. Among the gene functions highly associated with microbes, chloroplast-related functions showed the greatest enrichment. This is consistent with reports linking chloroplasts to plant defense and pathogen invasion as well as photosynthesis (Lu and Yao, 2018). This non-self host-response profile we describe is less immune-centered than that recently described for the non-self *A. thaliana* response (Maier et al., 2021). This difference is unsurprising given that (i) our study examined outdoor-grown plants instead of plants infected with individual microbes in a controlled environment, (ii) we profiled host expression at a very late stage of the host-microbiota interaction (after a few months of growth) instead of just nine days post-infection, and (iii) we describe the host response at the micrometer scale in different regions of individual leaves rather than the average response among homogenized leaves. Despite these methodological and conceptual differences, both studies revealed some similarities, such as the association between microbial infection and the immune-related genes AT1G02930 (GSTF6), which was among the 24 general non-self response genes that were discovered.

In conclusion, the versatility of SmT bodes well for its potential application to the many other tissue types ranging from plants to animals, including humans, where local differences in microbial colonization are an important determinant of health or disease.

## Methods

### Bacterial leaf-infiltration assay for microscopy

Seeds of *Arabidopsis thaliana* (accession Col-0) were surface sterilized by an overnight incubation at -80°C followed by washing with ethanol (5-15 min shaking in a solution of 75% EtOH [Sigma-Aldrich, USA] and 0.5% Triton X-100 [Sigma-Aldrich, USA], followed by a 95% EtOH wash and drying in a laminar flow hood). Stratification was done in a 0.1% agar solution at 4°C for 7 days before planting. Seeds were sown on potting soil (CLT Topferde; www.einheitserde.de), in 60-pots trays (Herkuplast Kubern, Germany). During the first two days after sowing (the germination period), the trays were covered with a transparent lid to reduce the likelihood of pest infection. Indoor growing conditions were as follows: Cool White Deluxe fluorescent bulbs (25 to 175 μmol m^-2^ s^-1^), 23°C, and 65% relative humidity. Plants were grown under long day conditions (16 h of light) for 15 days before syringe-infiltration with mCherry-tagged *Pseudomonas syringae pv. tomato* DC3000 (Pst DC3000) at OD_600_=0.001. Only half of the leaf was infiltrated (in relation to the main vein) - as visualized by the extent of soaking - so that the uninfected part could be used as a control. An 3xmCherry construct had been inserted at the *attn7* site and was a kind gift of Brian Kvitko.

Pst DC3000 was grown overnight in Luria Broth with the appropriate antibiotics (gentamicin and nitrofurantoin, 5 μg/mL each), then diluted 1:10 on the following morning, and was grown for an additional 4 hours to initiate the log phase, after which the bacteria were centrifuged at 3500 g for 90 seconds, and resuspended in 10 mM MgSO_4_.

Three days after infections, leaves were dissected and placed on 0.5x MS medium with agar (Duchefa, M0255), inspected under a Zeiss Axio Zoom.v16 fluorescence stereomicroscope to verify that the mCherry signal was present, and immediately flash-frozen in liquid N_2_. The leaves were stored at -80 °C before cryosectioning.

### Imaging of bacterial infected leaves

Infected *A. thaliana* leaves were imaged on a Zeiss Axio Zoom.v16 fluorescence stereomicroscope, equipped with an LED array for transmitted illumination and an X-Cite XYLIS LED from Excelitas Technologies for epi-illumination. All leaves were imaged using a PlanNeoFluar Z 1x/0.25 dry objective and a Hamamatsu ORCA-Flash4.0 digital CMOS camera (c11440-22C) with 2x2 binning. mCherry-tagged Pst DC3000 was detected using the Zeiss filter set 45 (00000-1114-462), which includes a 560/40nm excitation filter, a 630/75 nm emission filter, and a 585 nm dichroic filter. Brightfield images were acquired as references for the outline of the leaves for the analysis. The camera exposure time and light intensity were 220 ms and 5%, respectively, for the mCherry-tagged Pst DC3000. The images of infected leaves have a pixel size of 18.6 µm^2^ and were acquired at a 7x magnification. Image acquisition was done using the ZEN 2.1 software package.

### Outdoor-grown plants

For the analysis of microbial hot spots, microbial interactions and host responses to wild microbiomes, seeds of *Arabidopsis thaliana* (Accession Col-0) were germinated and grown indoors for seven short days (8 h of light). On 27 February 2019, the trays were placed outdoors near the Max Planck Institute for Biology Tübingen in a naturalized environment surrounded by other plants. Plants were irrigated weekly with regular tap water. Twenty-seven days after outdoor planting, individual leaves were sampled and immediately flash-frozen in liquid N_2_. Leaves from different plants were stored separately at -80 °C before cryosectioning.

### Multimodal array structure

Spatial metaTranscriptomics uses multimodal slides (10x Genomics Inc.) with capture areas of 6.5 x 6.5 mm. Each capture area comprises 4,992 spots, with diameters of 55 μm each. The spots are covered with capture probes in the following proportion: 45% 16S rRNA probes, 45% 18S rRNA/ITS probes and 10% poly-d(T) probes.

### Probe design

Probes were designed using two approaches: one based on established primers of the relevant marker genes (P799 (Hanshew et al., 2013) and P902 (Hodkinson and Lutzoni, 2009)) and a *de novo* approach (P1265 and P479) (**Supplementary Figure S11**). On average, the probing sites were 100 nt upstream of the target site. In general, we aimed to maximize two variables: the conservation of the probe sites and the variability of the 100 nt downstream target sites. The *de novo* design process was adopted because previously designed primers were suboptimal with respect to these criteria.

Previously designed primers were used as templates due to their wide usage in the field, which is indicative of useful specificity – it implies that they have a wide taxonomic range and good ability to exclude host reads such as those originating from 16S chloroplast rRNA. Four probes were designed based on previous primers; the 16S probes 16S:P799 and 16S:P902 were based on the mainstream primers 799F (Hanshew et al., 2013) and 902R (Hodkinson and Lutzoni, 2009), respectively. Additionally, the eukaryotic capture probes probes 18S:P-ITS1 and ITS:P-ITS7 were based on the mainstream primers ITS1F (Gardes and Bruns, 1993) and ITS7F (Ihrmark et al., 2012), respectively. To fit the primers to the annealing conditions of the array, we reversed-complemented all forward-oriented primers (i.e. all of them but 902R; the target RNA is single stranded, so reversal of the primer orientation was needed to capture it) and elongated them to obtain 35-45 bp long sequences, as recommended for microbial profiling in microarray systems (Gardner et al., 2013). To this end, 16S rRNA and ITS custom databases were downloaded (on 29 April 2020) from NCBI GenBank and the sequences downstream of the primer (up to 100 nt, including the primer) were extracted. These sequences were then aligned using the software Clustal Omega (v1.2.4) and the sequence profiles were plotted using weblogo (v3.7.5). The primers were elongated by manual inspection of the resulting weblogo. The length and degeneracy level were limited to obtain fewer than 35,000 unique probe sequences.

In addition to these probes, two *de novo* 16S probes were designed to complement the primer-based probes (as shown in **Supplementary Figure S11**) - 16S:P1265 (5’-GGT AAG GTT YYK CGC GTT GCD TCG AAT TAA ACC RCAT-3’) and 16S:P479 (5’-TCT CAG THC CAR TGT GGC YBD YCD YCC TCT CARR-3’). To design these probes, representative sequences were selected from the SILVA 16S database (v138.1) using CDHIT (v4.8.1) to the level of 99% sequence identity. Representative sequences were aligned using MAFFT (v7.245), and the sequence profile was plotted using weblogo (v3.7.5). In this process, we targeted highly variable regions with a conserved matching probing site.

### Sample preparation and sectioning

The leaves stored at -80°C were immersed in 50% Optimal Cutting Compound (OCT, Sakura) in PBS (Medicago). Embedded samples were frozen in a cryotome (Cryostar NX70, ThermoFisher Scientific) and sectioned to obtain 14-μm longitudinal sections. Tissue sections were then laid over the multimodal capture areas of the arrays.

### Tissue optimization experiment

Tissue permeabilization conditions were identified using a modified variant of a previously reported protocol (Giacomello and Lundeberg, 2018). Briefly, after attaching of the tissue section to the slide surface containing 100% poly-d(T) capture probes the tissue was fixed in methanol (VWR) at -20°C for 40 minutes and stained with 0.05% Toluidine Blue (Sigma-Aldrich) at room temperature for 2 minutes. Tissue sections were imaged using a Zeiss AxioImager 2T and a Metafer slide scanning system (v. 3.14.2, Metasystems). They were then permeabilized with pepsin (Sigma-Aldrich) in 0.1%/0.1 M hydrochloric acid (Fluka) at 37 °C for 30 min. The plant mRNA molecules that had hybridized to the capture probes were reverse transcribed to cDNA using SuperScript III (Invitrogen, ThermoFisher Scientific) and Cy3-dCTP-nucleotides (PerkinElmer) at 42°C overnight. Tissue sections were removed from the slide surface by incubation for one hour at 37°C in a hydrolytic enzyme mixture consisting of pectate lyase (Megazyme), xyloglucanase (Megazyme), xylanase 10A (Nzytech), β-mannanase 26A (Nzytech), and cellulase (Worthington) in monobasic sodium citrate (Sigma-Aldrich), pH 6.6. They were then incubated with 2% β-mercaptoethanol (Calbiochem) in RLT buffer (Qiagen) and proteinase K (Qiagen) in PKD buffer (Qiagen) for one hour each. Finally the fluorescent cDNA footprint was imaged using an Innoscan 910 (Innopsys) slide scanning system and Mapix image analysis software (v. 9.1.0, Innopsys) with a pixel size of 5.0 and a gain of 50.

### Sequencing library preparation

Sequencing libraries were prepared according to the Visium protocol (10x Genomics Inc.) with the following modifications: multimodal slides with leaf sections attached to the capture areas were incubated for 2 minutes at 37°C followed by a 40-minute fixation in methanol (VWR) at -20°C. Capture areas were washed with PBS (Medicago) and incubated for 2 minutes at 37°C. Tissue sections were stained for 2 minutes with 0.05% Toluidine Blue (Sigma-Aldrich) at room temperature followed by two washes with ultrapure water and warming at 37°C for 2 minutes. The slides were mounted with 85% glycerol (Merck) and the bright-field images were acquired with a Zeiss AxioImager 2X microscope and a Metafer slide scanning system (v. 3.14.2, Metasystems) at 20x magnification. To increase permeabilization efficiency and reduce the effect of secondary metabolites, the slides were incubated in 2% (wt/vol) polyvinylpyrrolidone 40 (PVP-40, Sigma-Aldrich) at room temperature for 10 minutes. Host-plant and eukaryotic microbial cells were permeabilized using the permeabilization enzyme (10x Genomics Inc.) at 37°C for 30 minutes. Bacterial organisms were permeabilized using 10 mg/ml lysozyme from chicken egg white (Sigma-Aldrich) in 0.05 M EDTA pH 8.0 (Invitrogen), 0.1 M Tris-HCl, pH 7.0 (Invitrogen) for 30 minutes at 37°C.

The rest of the SmT workflow followed the procedure described in the Visium Spatial Gene Expression user guide with the following modification: reverse transcription was performed using 2% (wt/vol) PVP-40 instead of nuclease-free water to reduce adverse impacts due to secondary metabolites and cDNA was amplified by performing 12-15 PCR cycles. Libraries were sequenced using Illumina Nextseq 2000 and Nextseq 1000/2000 P2 or P3 Reagents (200 cycles) kit.

### Preprocessing of the reads and bright field images

Template switch oligo and long poly-A stretches were removed from Read 2 using cutadapt (v. 2.9, Martin, 2011). The location of the tissue was determined using the Loupe Browser (v. 5.1.0, 10x Genomics) in which all the spots containing at least 25% of the tissue were selected and their locations (i.e., x and y coordinates) were recorded.

### Read alignment

TSO- and poly-A trimmed reads were analyzed using the ST Pipeline (v. 1.7.9, Navarro et al., 2017), which enables simultaneous analysis of the spatial location, unique molecular identifier, and mRNA molecule. First, the pipeline trimmed poly-N stretches which are longer than 15 bp. Read 2 was then mapped against the *A. thaliana* TAIR10 genome release (Berardini et al., 2015) using the STAR (v. 2.7.7a, Dobin et al., 2013) mapping tool and annotated with htseq-count 1.0 (Anders et al., 2015). The spatial barcode in Read 1 was demultiplexed using Taggd (Costea et al., 2013) and the information from Read 1 and Read 2 was combined. The ST Pipeline then grouped the reads based on the spatial barcode, gene and genomic location. Finally, the unique molecules were identified using a unique molecular identifier and the counts were compiled into the gene-count matrix.

### Taxonomic assignment of microbial reads

Reads were mapped against the *A. thaliana* reference genome using STAR (v. 2.7.7a, Dobin et al., 2013) and all reads aligning to the genome were discarded, leaving putative microbial reads. Next, read datasets were demultiplexed based on their probe types (i.e 16S rRNA, ITS/18S rRNA). For each probe dataset, the reads were first clustered into representative sequences by the fastx_uniques module of usearch (v. 11.0.667, Edgar, 2010). Next, the representative sequences (query) were searched for the best homolog (hit) in the NCBI NT database [downloaded January 2021] (Sayers et al., 2021) using MMseqs2 (v. 1f30213, Mirdita et al., 2019; Steinegger and Söding, 2017). For each query, all of the best hits (i.e., those with the highest identical bit score and a taxonomic assignment on the genus level) were selected for further consideration. Next, the taxonomic assignment for a query was set as the lowest common ancestor (LCA) among the best hits as calculated by taxonkit (v. 0.7.2, Shen and Ren, 2021) using the NCBI Taxonomy database [downloaded on Jan 2021] (Schoch et al., 2020). For 18S rRNA/ITS probes, reads were further considered if they were classified as Eukaryota but not as unclassified, chloroplast, mitochondria, uncultured, *Streptophyta*, *Chordata*, or *Arthropoda* on the genus or the phylum levels. Similarly, for 16S rRNA probes, reads were considered if they were classified as Bacteria but not as unclassified, chloroplast, mitochondria, or uncultured on the genus level. Finally, reads were further filtered by their UMI, such that for each spatial location, only one representative read with a given UMI was retained. The number of reads considered for each dataset is provided as **Supplementary Table S8**.

### Pst DC3000 infection experiment - data processing

Processed, aligned reads were analyzed using STUtility (v. 0.1.0, Bergenstråhle et al., 2020). To exclude low quality spots, the *A. thaliana* host data and bacterial unique molecules were summed together and every spot with fewer than 20 unique molecules was discarded. In addition, each spot contained at least 10 unique genes/taxa. The visualized genes and taxa were log10 normalized and projected on a bright field image of the tissue section with an opacity of 0.75.

### Enrichment experiment

Glass slides bearing a multimodal capture array (10% poly-d(T) probes, 45% bacterial 16S rRNA probes and 45% eukaryotic 18S rRNA/ITS probes), a 100% poly-d(T) array, a 100% bacterial 16S rRNA array, and a 100% eukaryotic 18S rRNA/ITS array were used. Three leaves were sectioned on each of these capture slides meaning every leaf had a consecutive section on each array type. Sequencing libraries were prepared as per the above protocol and sequenced with Nextseq 2000 (Illumina). The reads were annotated as described above and analyzed using R (v. 4.0.5).

STUtility (v. 0.1.0) was used to read the *A. thaliana* data to an object and sums of gene values were log_10_-transformed. Pairwise Pearson correlation coefficients were calculated and visualized with the corrplot package (v. 0.92) function corrplot.mixed using significance levels of 0.001, 0.01 and 0.05, with hierarchical clustering permitted.

For bacterial 16S rRNA and eukaryotic 18S rRNA/ITS data, unique molecules were summed together per taxon, generating a table containing the sum of unique molecules, phylogenetic paths, and metadata relating to section identification. Any annotations to phylum *Streptophyta* were removed, after which the data were divided into bacterial and fungal datasets based on their superkingdom. Pairwise correlations were only calculated for classified reads. We performed the analysis three times – first with all taxa and then with only the most highly expressed 500 and 20 taxa.

### Validation of SmT with amplicon sequencing

To compare the performance of SmT to that achieved with amplicon-sequencing, seeds of *A. thaliana* (accession Est-1) were surface-sterilized and stratified at 4°C for one week in a refrigerator, and then sown in plastic trays (Herkuplast, Ering, Germany) filled with wild soil from the Heuberger Tor experimental site of the University of Tübingen (Germany). The seeds were left outside to germinate in the same field in late September. The plants developed and overwintered without supplemental watering. Additional plants in each pot were thinned in January 2020 with tweezers, and individual plants were sampled at the end of March 2020 before flowering. The sampling protocol involved cutting the mature rosettes with sterile scissors, placing them in sterile 50 mL centrifuge tubes, and vigorously shaking them in sterile water. The water was then dumped and new water was added until the leaves released no further dirt. After washing, plants were immediately flash-frozen in liquid N_2_, and subsequently stored at -80°C prior to nucleic acid extraction. Both DNA and RNA were extracted from each plant. The entire rosette was lysed in a buffer containing 2% β-Mercaptoethanol to extract all nucleic acids while preserving RNA. One proportion of the lysate was used for RNA extraction by the phenol/chloroform protocol, while another portion was used to extract DNA following a previously described potassium acetate and SPRI bead protocol (Regalado et al., 2020). The DNA moiety was used for 16S rDNA amplicon sequencing. Two sets of primers were used: (i) 515F-806R (V3-V4) in combination with plastid-blocking clamps (Lundberg et al., 2013), and (ii) 799F-1192R (V4-V6), which does not amplify chloroplasts and for which the mitochondrial amplicon was removed by gel extraction (Bulgarelli et al., 2015). The RNA moiety was used for SmT, using the same pipeline as for all other samples with the exception that crude extracts were used in place of leaf samples (so spatial information was not extracted). A total of 300 μg of RNA was used for the array. In total, four plant samples were used for 16S rRNA profiling, comparing two amplicon sequencing primer sets to the SmT array. The reads obtained by amplicon sequencing were analyzed in the same way as the array reads, excluding the initial mapping to the *A. thaliana* TAIR10 database. For both of the methods, the reads were subsampled to the same sequencing depth. See the ‘Read alignment’ and ‘Taxonomic assignment of microbial reads’ sections for information about the full pipeline.

### Analysis of microbial hotspots

Microbial hotspots (based on 16S rRNA/ITS reads) were identified using the Getis-Ord G statistic (Ord and Getis, 1995) as implemented in the localG function of the R spdep package (v. 1.1.11, Bivand and Wong, 2018). The calculation was performed using a 2x2 grid applied to the count matrix resulting from the sum of reads belonging to the 50 most abundant genera (separately for 16S rRNA/ITS reads). The p.adjustSP function of the R spdep package was used with the BH-FDR (Benjamini and Hochberg, 1995) method to correct the G stats p-values while accounting for the number of neighbors of each region. Hotspot spatial maps were plotted using the R tmap package (v. 3.3-2).

### Microbial interaction network analysis

Microbial interactions were inferred based on the Spearman rank correlation coefficient (SRCC) values of the reads count associated with each pair of genera. Specifically, for each pair of microbial genera, in each leaf section, SRCC was calculated accounting for all spots of the array (i.e., each spot on the array was considered as a ‘sample’ for each genus). We considered pairs of genera to be interacting if their SRCC corrected p-value (BH-FDR) was below 0.05. Next, to account also for the spatial organization of microbes in the array, we computed the SRCC value of each candidate pair based on shuffled abundance matrices. This step, repeated 1,000 times, results in an empirical null distribution of expected SRCC values where the spatial association between a paired genera is random. The shuffled count matrix was generated by using the permatfull function implemented in the R vegan package (v. 2.5.6) while keeping the total number of reads associated with each genus across all samples (spots) constant (i.e. argument fixedmar = "columns"). Finally, the significance of each candidate pair of genera was calculated by comparing the SRCC value based on the unshuffled count matrix to the empirical null pair distribution (North et al., 2002) following an BH-FDR correction. Microbial interactions were considered to be also spatially significant if their corrected-empirical p-value was below 0.05. The network was created based on these microbial pairwise correlations values using the R igraph package (v. 1.2.6) and plotted using the R ggraph package (v. 2.0.5).

### Host mRNA clustering

For the *A. thaliana* host data, the counts were filtered using STUtility (v. 0.1.0, Bergenstråhle et al., 2020) by removing the low quality spots and genes containing at least 10 and 30 counts, respectively. In addition, each spot was required to have at least 10 genes and each gene was required to cover at least 20 spots. Chloroplast, mitochondrial, ribosomal and non-coding genes were filtered from the data set because many of them are not polyadenylated and might contain genes captured with 16S rRNA and 18S rRNA/ITS probes. Finally, after the filtering steps, the spots with fewer than 10 genes were removed because they were considered to be of low quality.

Each section was normalized individually using the Seurat (v. 4.1.0, Hao et al., 2021) function sctransform (Hafemeister and Satija, 2019) to eliminate intra-section batch effects. To reintegrate the sections back together, anchor features were selected and the whole data was scaled based on these features. Principal component analysis (PCA) was performed on this data using identified variable features. Based on the results of the PCA, the inter-section batch effects (experiment date, plant and leaf) were removed with Harmony (v.0.1.0, Korsunsky et al., 2019) using a diversity clustering penalty of 4 and PCA dimensions of 1 to 8.

Normalized gene counts were projected onto 2D leaf section images using UMAP (McInnes et al., 2018) with the eight first dimensions from Harmony and a resolution of 0.22. To identify cluster specific markers, raw counts were normalized using the NormalizeData function with LogNormalize as a normalization method and the FindAllMarkers function with the parameters of test.use="poisson" and logfc.threshold=0.15.

### Host-response analyses

We used the Boruta algorithm (Kursa et al., 2010) to determine which set of *A. thaliana* genes is important to explain the microbial load on each spot of the array. Briefly, we modeled the relationship between the expression profile of all A. thaliana genes - *G_1_…G_n_* and *M* - the sum of the 50 most abundant Bacterial/Fungi genera in each spot of the array (M): M ∼ G_1_…G_n_. We treated the task as a regression problem and used the random forest algorithm (Breiman, 2001) to calculate the importance of each gene in the model. Next, we used Boruta to assign a significance score for each gene based on its importance for the model’s accuracy. For this purpose we used the R implementation of the Boruta package (v. 7.0.0) with 1000 trees. This procedure was performed for each leaf-section, once using the un-normalized read counts and once using the Getis-Ord G statistic value, treating each spot as an observation. Overall, a gene was considered further if it was found to be significant by Boruta for at least one measure (i.e., reads count or G statistics), and if its SRCC p-value (after FDR correction) was below 0.01. GO enrichment analyses were performed with the DAVID webserver with the DAVID knowledgebase v2022q1 (Huang et al., 2009; Sherman et al., 2022).

### Data and code availability

Sequencing data have been deposited at NCBI-SRA under the Bioproject PRJNA784452 and will be made publicly available upon publication. Tissue images will be available at the SciLifeLab data repository (https://www.scilifelab.se/data/repository) upon publication. Scripts written for the analyses described in this paper are available on Github (https://github.com/giacomellolab/SpatialMetaTranscriptomics).

## Acknowledgements

We thank Uppsala Multidisciplinary Center for Advanced Computational Science (UPPMAX) for providing computational infrastructure and the Bio-Optics Facility of the Max Planck Institute for Biology Tübingen for assistance with microscopy. We thank 10X Genomics Inc. for providing multimodal arrays. The authors thank Ludvig Larsson for discussions on data analysis. S.G. was supported by Formas grant 2017-01066 and VR grant 2020-04864. H.A. was supported by a fellowship from the Alexander von Humboldt Foundation. Supported by funds from Formas and VR to S.G. and the Max Planck Society to D.W.

## Author contributions

Conceptualization: S.G. and D.W.

Methodology: S.G. designed SmT and relative experiments; S.S. and O.S. contributed to the design of SmT; O.S. designed Pst DC3000 experiments and sample collection; H.A. developed the hotspot and network analysis workflow.

Investigation: S.S. performed SmT experiments; O.S. inoculated *A. thaliana* leaves with Pst DC3000 and collected wild Arabidopsis leaves; V.d.O.C. performed leaf fluorescence imaging; D.S.L. generated amp-seq data; S.S., O.S., H.A., D.W. and S.G. interpreted the results.

Formal analysis: S.S. performed read alignments and unsupervised clustering of the host data;

O.S. and S.S. analyzed the enrichment level of the different probe percentages; H.A. and O.S. designed bacterial and fungal capture probes. H.A. performed taxonomical annotation of the sequencing data, calculated microbial hotspots and spatial networks, performed GO and Boruta analyses.

Validation: S.S., O.S., H.A., D.W. and S.G.

Data curation: S.S. and H.A.

Writing – original draft: S.S., O.S., H.A., D.W. and S.G.

Writing – review & editing: All authors.

Visualization: S.S.

Supervision: S.G. and D.W.

Project administration: S.G., D.W., S.S. and O.S.

Funding acquisition: S.G. and D.W.

---

**Supplementary Figure S1:**
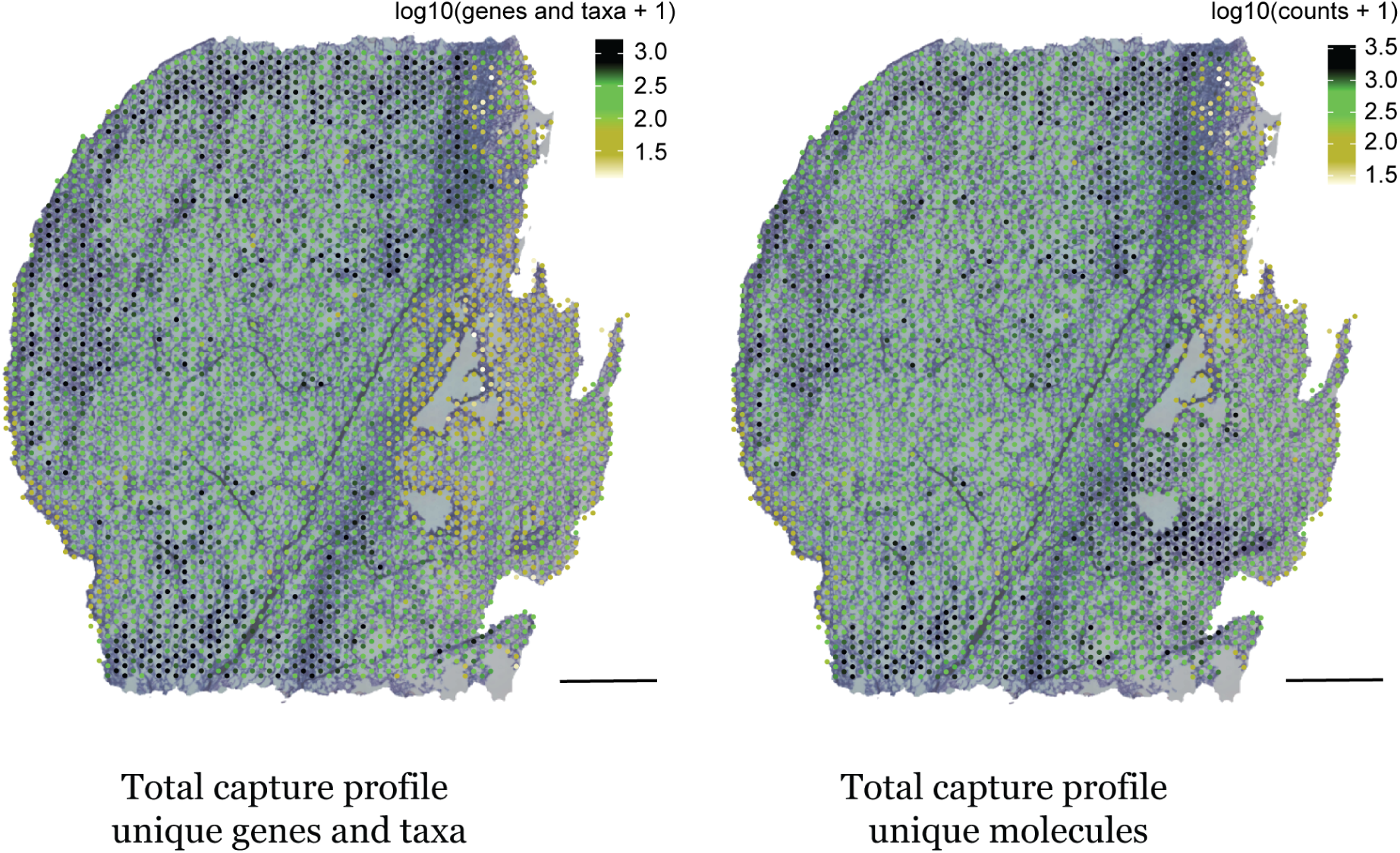
Capture profiles of unique genes and taxa and unique molecules across the tissue sections from *Pseudomonas* infected leaves. Scale is 1 mm.

**Supplementary Figure S2:**
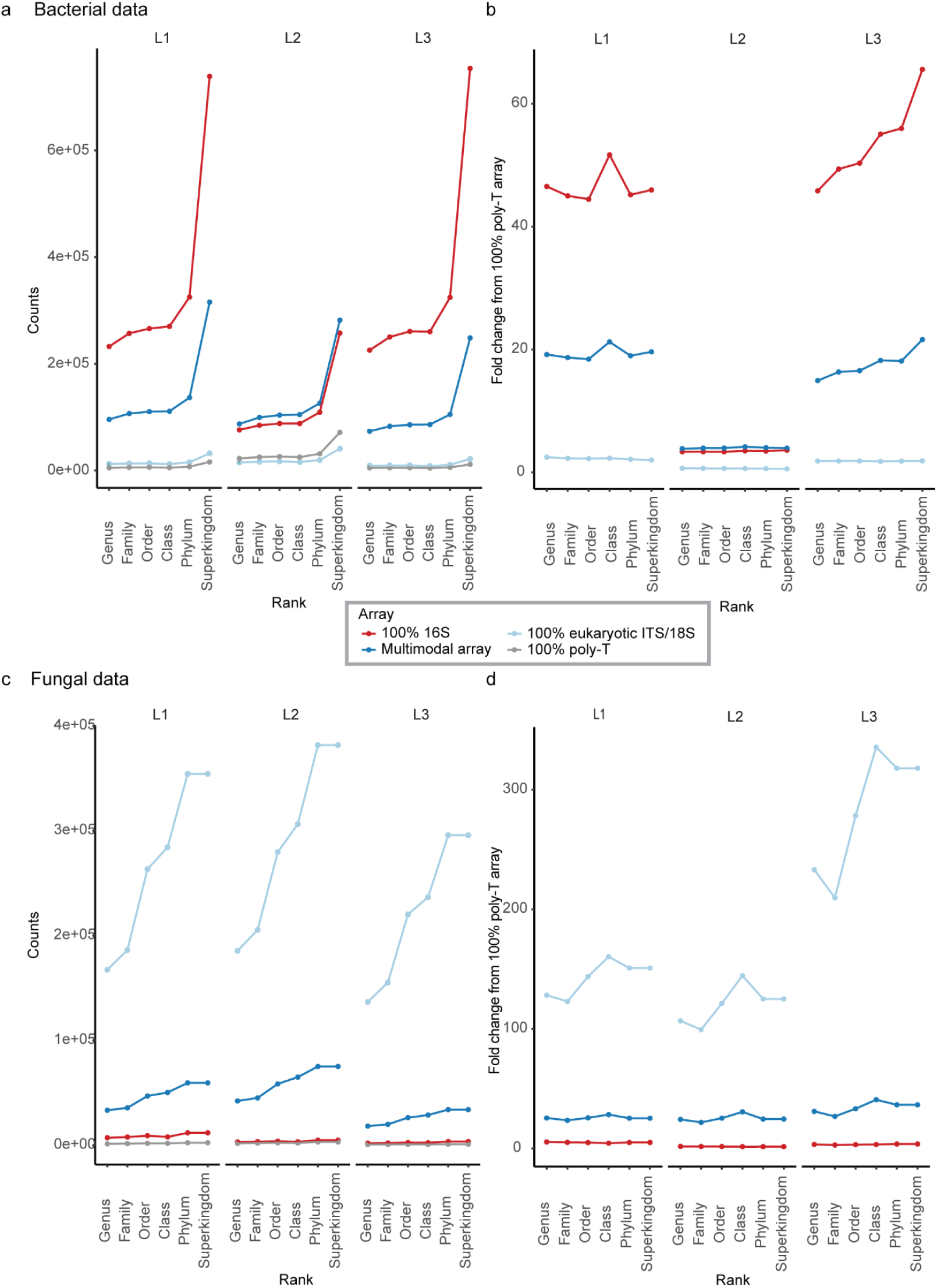
Number of unique molecules presented in each taxonomic level for bacterial data in each array (**a**) and the fold change between different taxonomic levels from unspecific binding where an array of 100% poly-d(T) probes has been used as a baseline (**b**). Fungal unique molecules and the fold change as a function of taxonomic level are presented in **c** and **d**, respectively. L1, L2 and L3 stands for Leaf 1, Leaf 2 and Leaf 3, respectively.

**Supplementary Figure S3:**
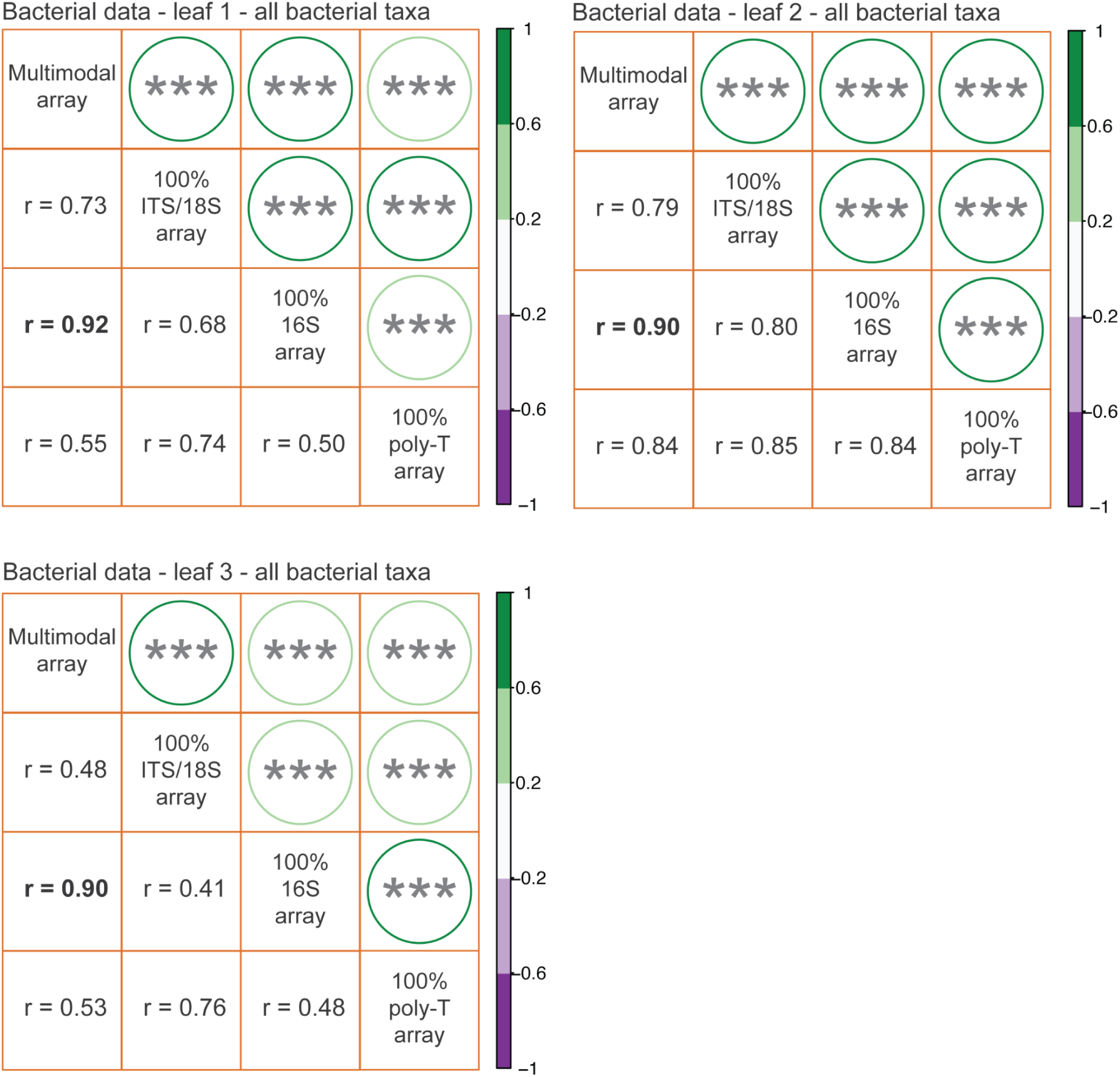
Pairwise correlations of bacterial 16S components between different array types in three leaves when the full bacterial profile with 1681 taxa is considered. *:P<0.05, **:P<0.01, ***:P<0.001. Correlation between the multimodal array and the 100% 16S array is in bold.

**Supplementary Figure S4:**
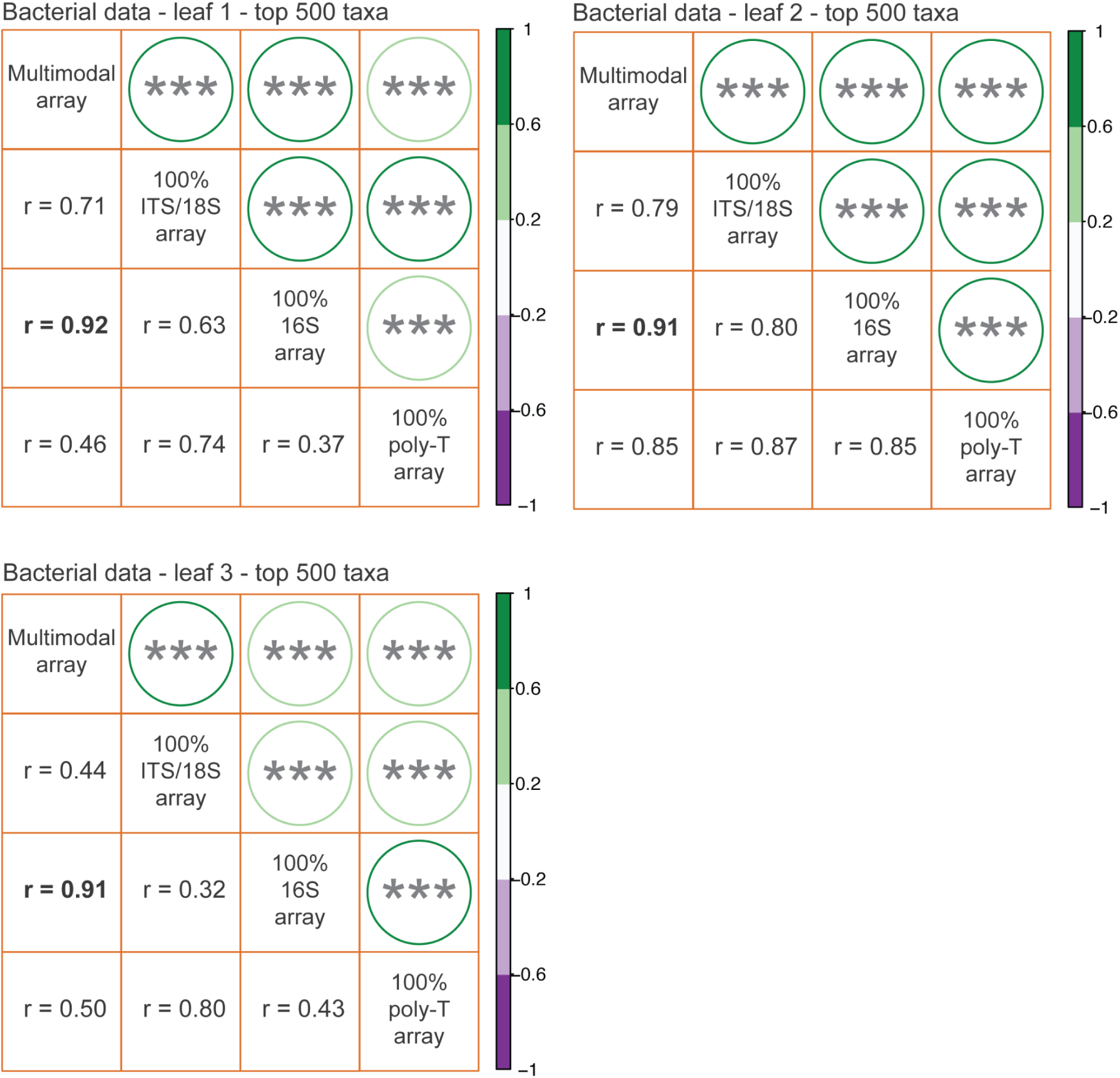
Pairwise correlations of bacterial 16S components between different array types in three leaves when the 500 most abundant bacterial taxa are considered. *:P<0.05, **:P<0.01, ***:P<0.001. Correlation between the multimodal array and the 100% 16S array is in bold.

**Supplementary Figure S5:**
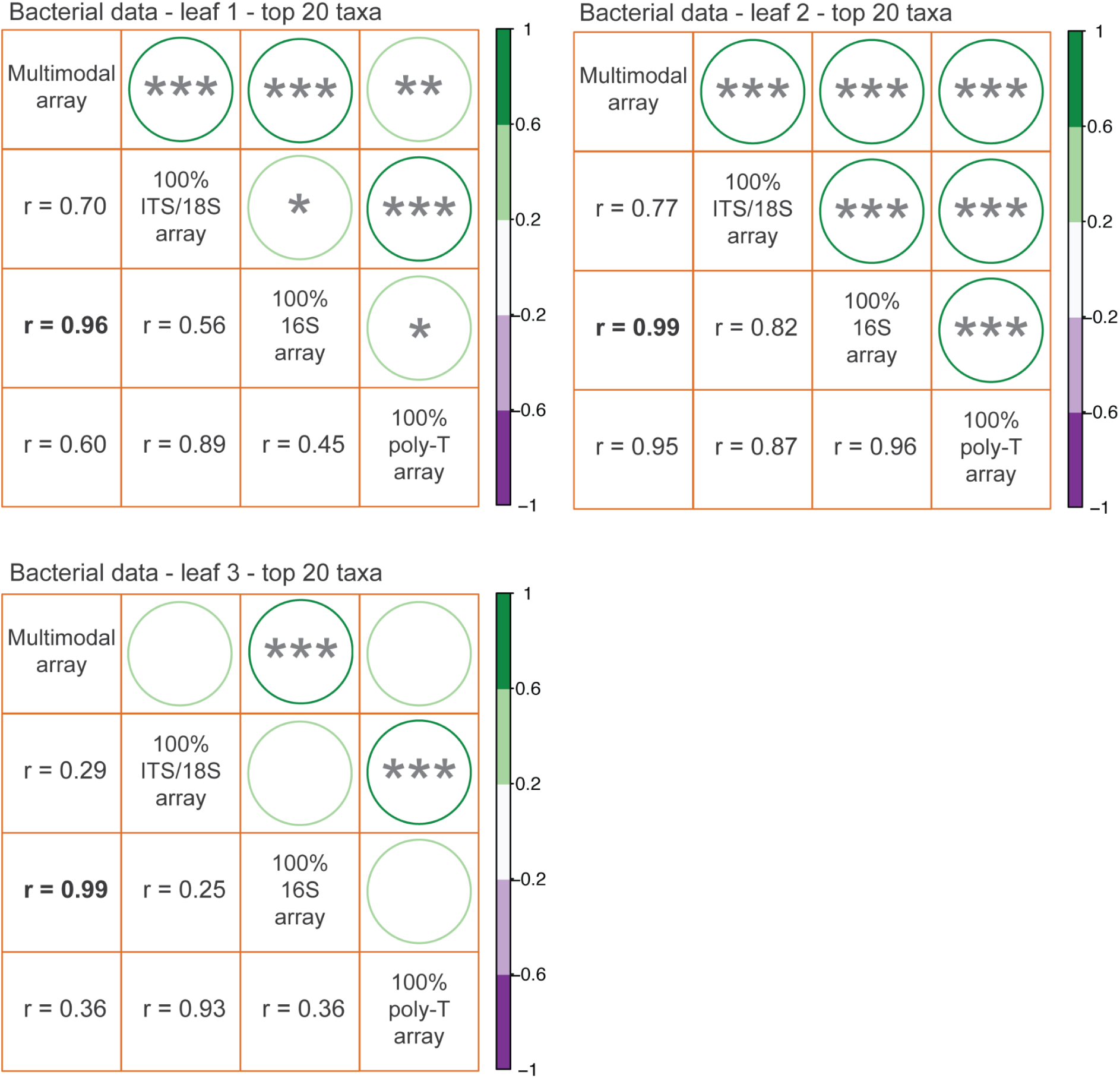
Pairwise correlations of bacterial 16S components between different array types in three leaves when the 20 most abundant bacterial taxa are considered. *:P<0.05, **:P<0.01, ***:P<0.001. Correlation between the multimodal array and the 100% 16S array is in bold.

**Supplementary Figure S6:**
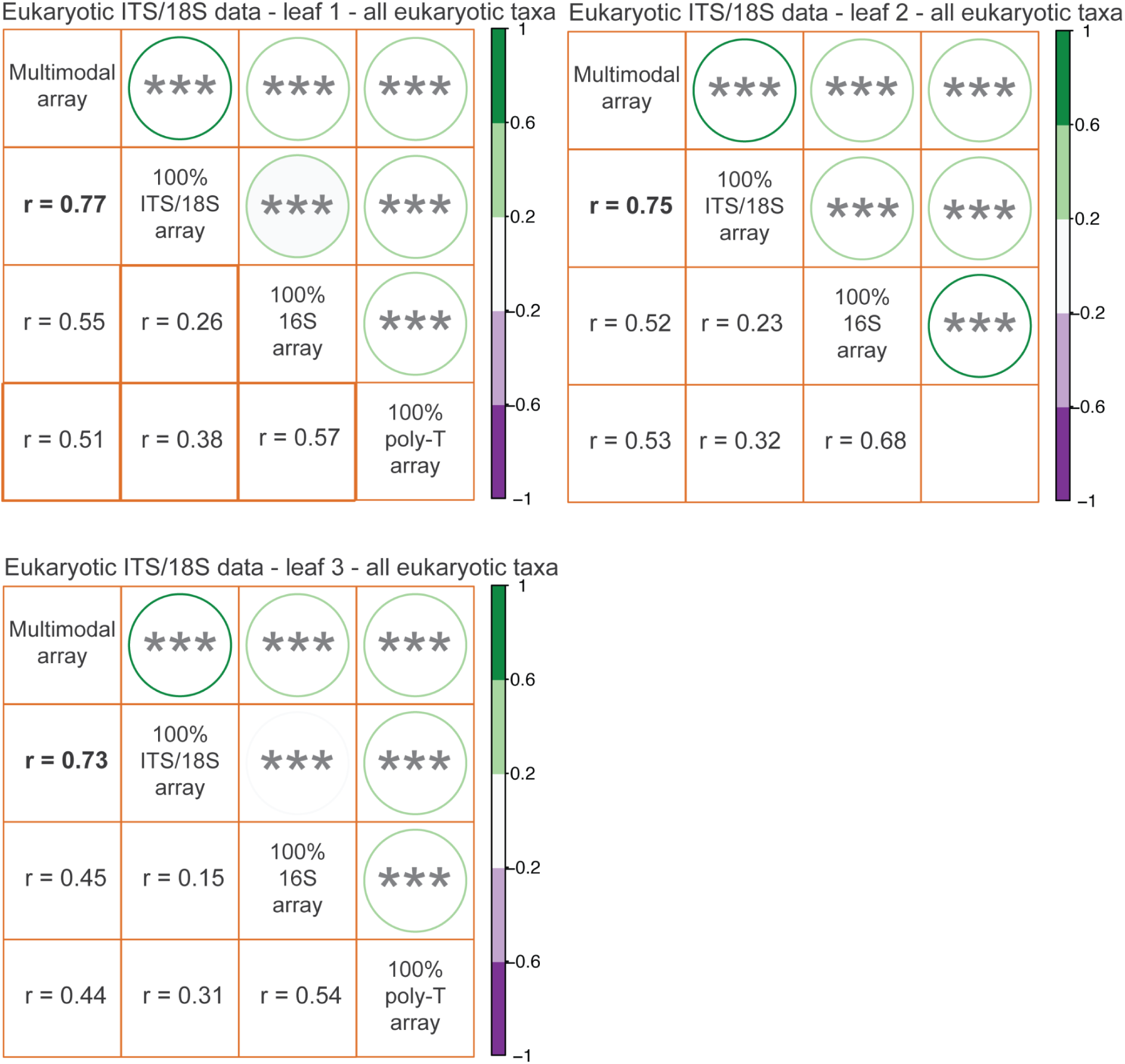
Pairwise correlations of eukaryotic ITS/18S components between different array types in three leaves when the full eukaryotic profile with 1660 taxa is considered. *:P<0.05, **:P<0.01, ***:P<0.001. Correlation between the multimodal array and the 100 % Eukaryotic ITS/18S array is in bold.

**Supplementary Figure S7:**
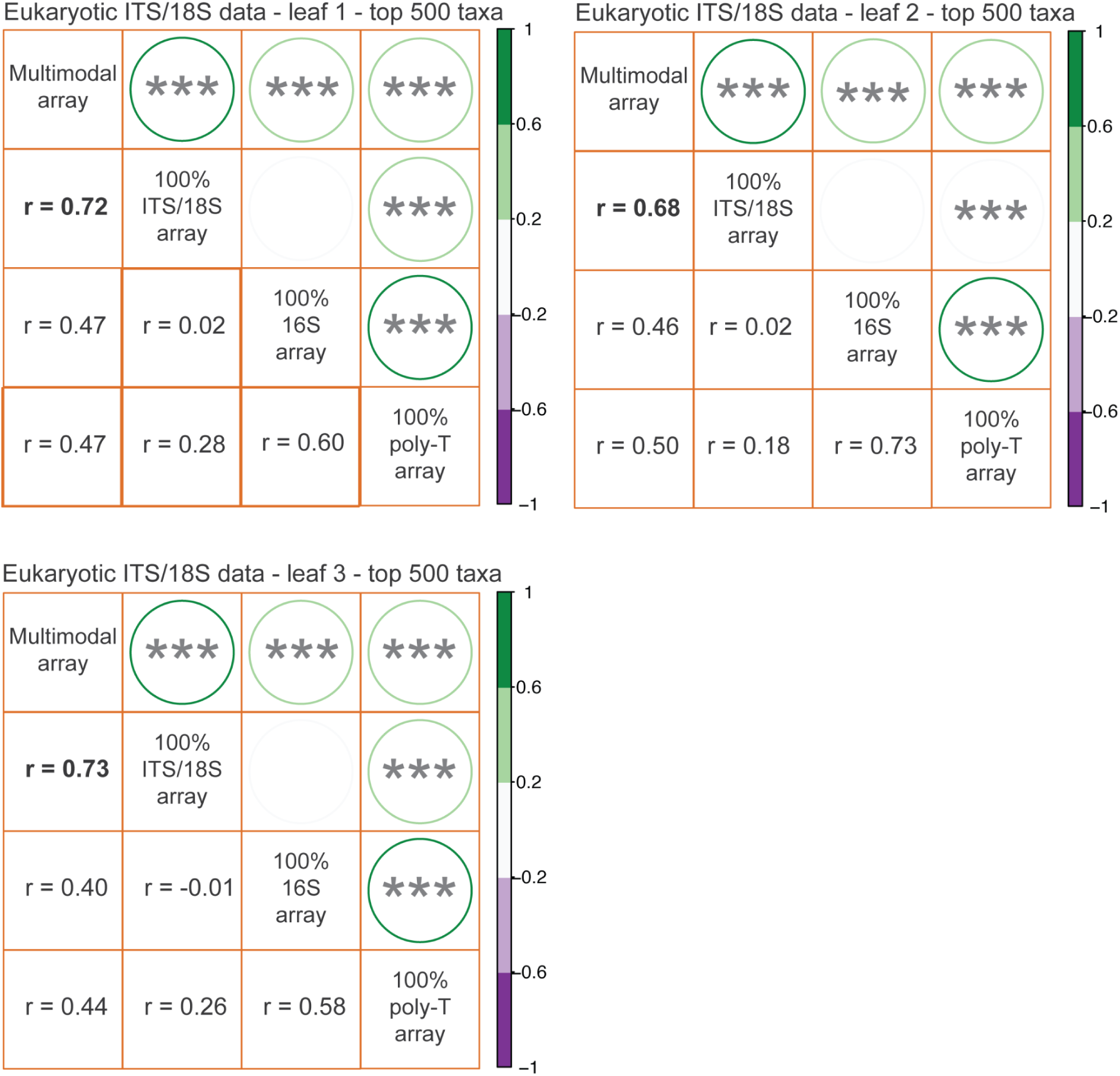
Pairwise correlations of eukaryotic ITS/18S components between different array types in three leaves when the 500 most abundant eukaryotic taxa are considered. *:P<0.05, **:P<0.01, ***:P<0.001. Correlation between the multimodal array and the 100% Eukaryotic ITS/18S array is in bold.

**Supplementary Figure S8:**
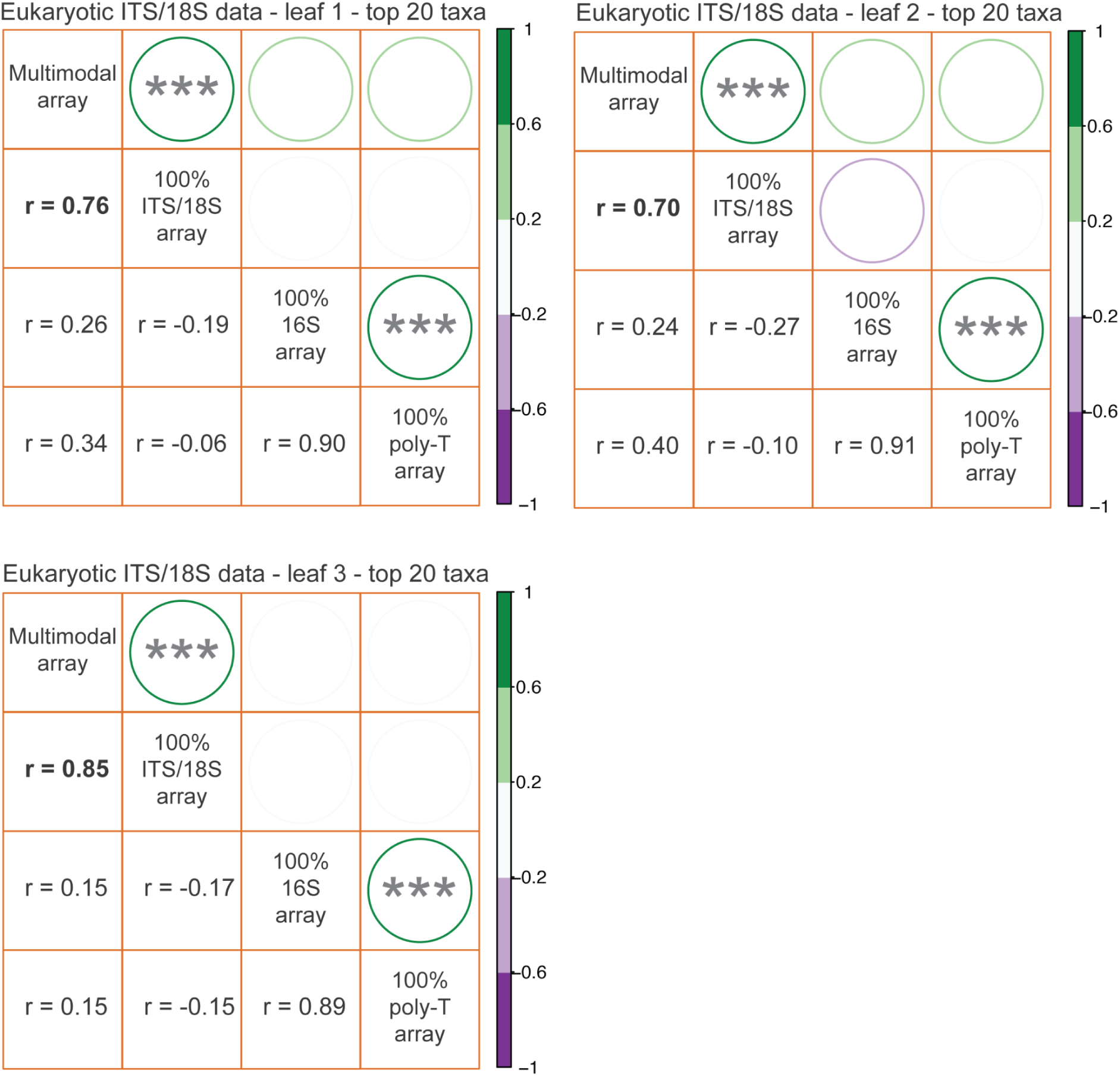
Pairwise correlations of eukaryotic ITS/18S component between different array types in three leaves when the 20 most abundant eukaryotic taxa are considered. *:P<0.05, **:P<0.01, ***:P<0.001. Correlation between the multimodal array and the 100% Eukaryotic ITS/18S array is in bold.

**Supplementary Figure S9:**
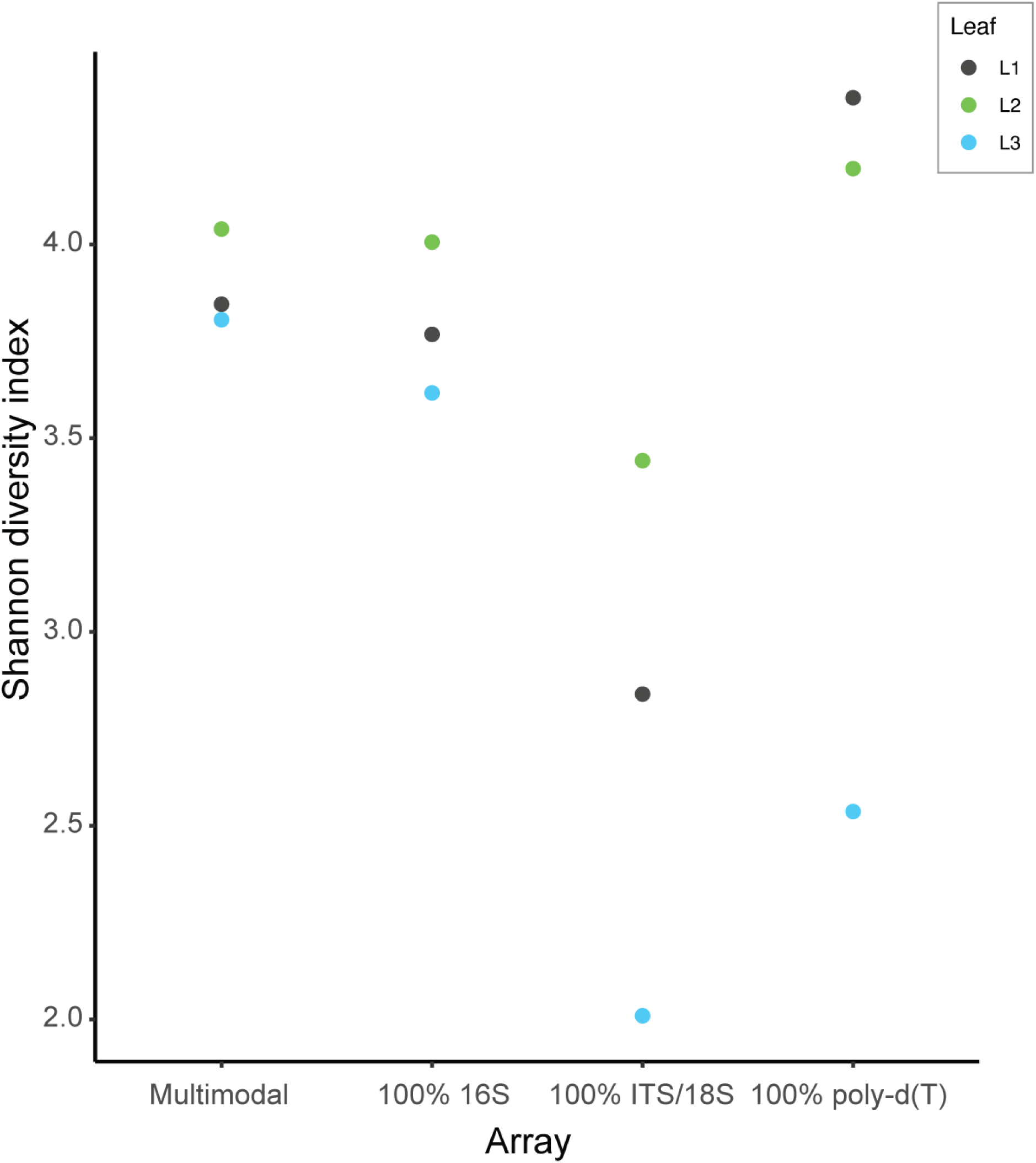
Shannon diversity index between different leaves in different arrays with bacterial assignment. L1, L2 and L3 stands for Leaf 1, Leaf 2 and Leaf 3, respectively.

**Supplementary Figure S10:**
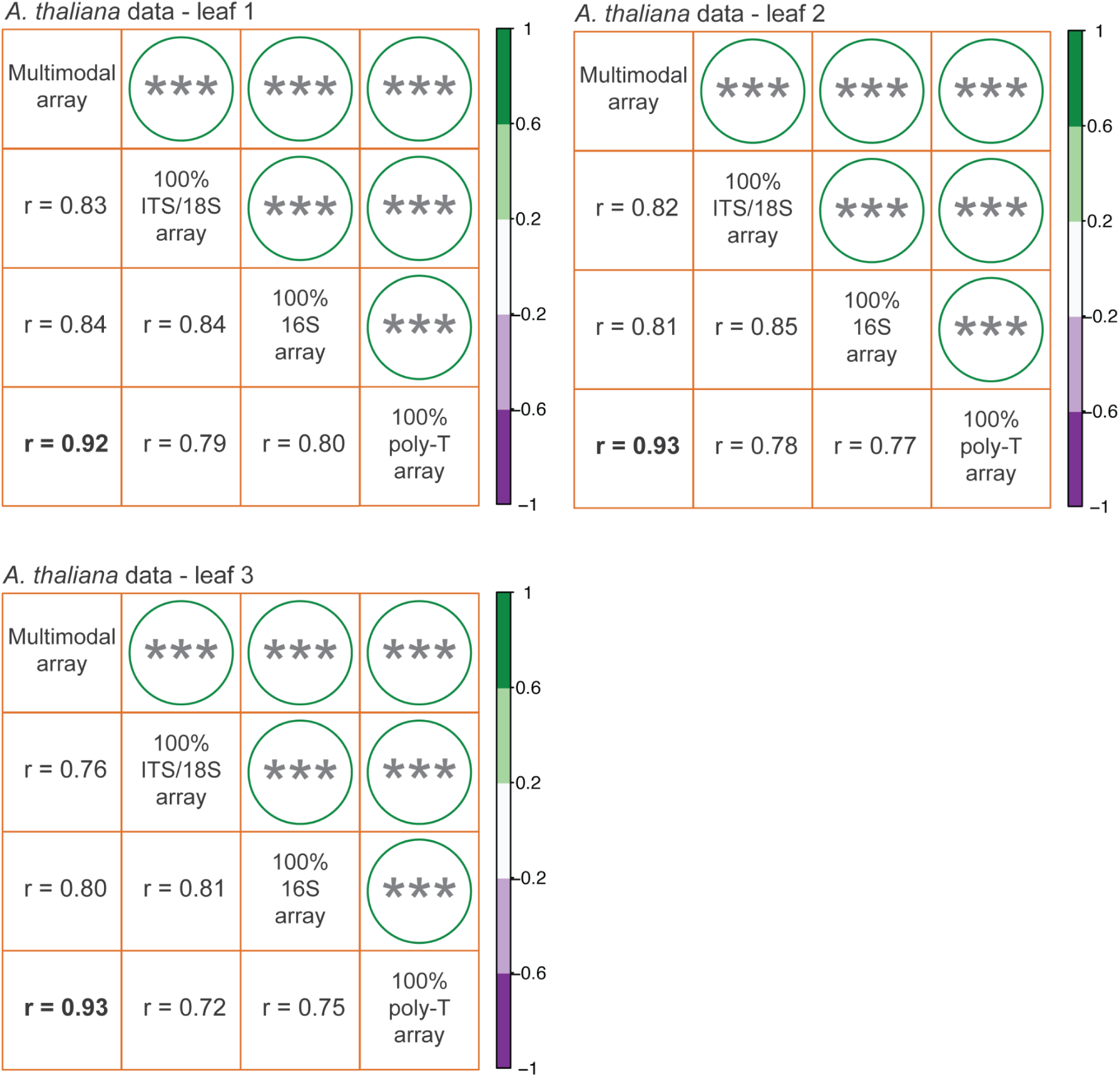
Pairwise correlations of the host *A. thaliana* component between different array types in three leaves. *:P<0.05, **:P<0.01, ***:P<0.001. Correlation between the multimodal array and the 100% poly-d(T) array is in bold.

**Supplementary Figure S11:**
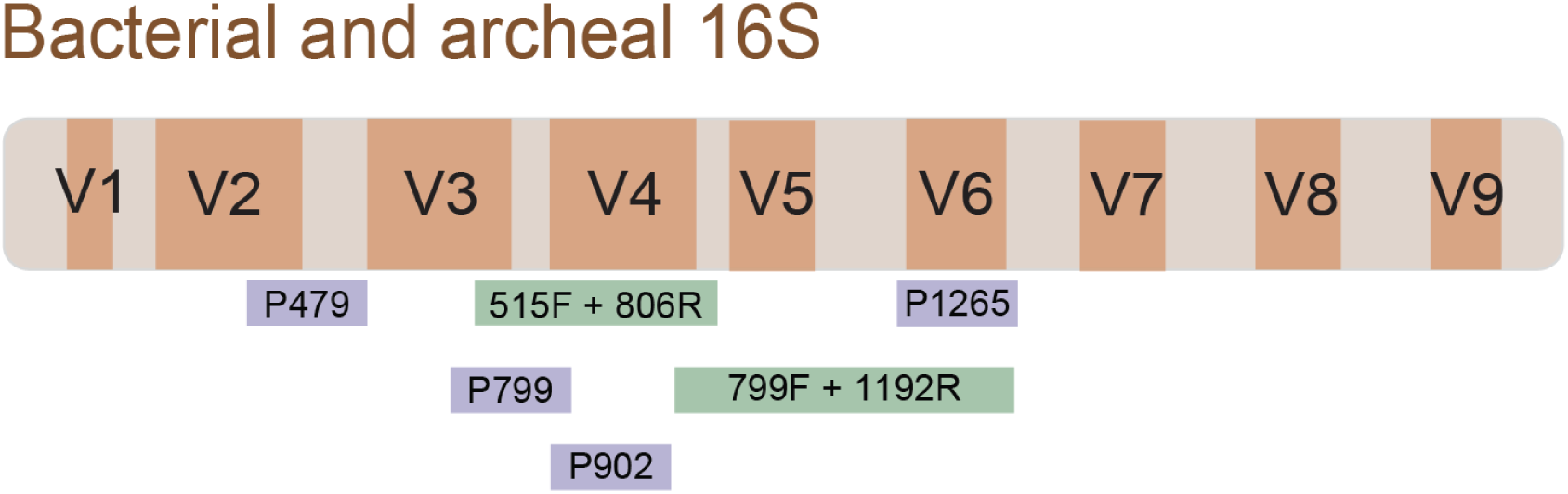
Bacterial and archeal target regions of the SmT probes (purple) and amp-seq primers (green).

**Supplementary Figure S12:**
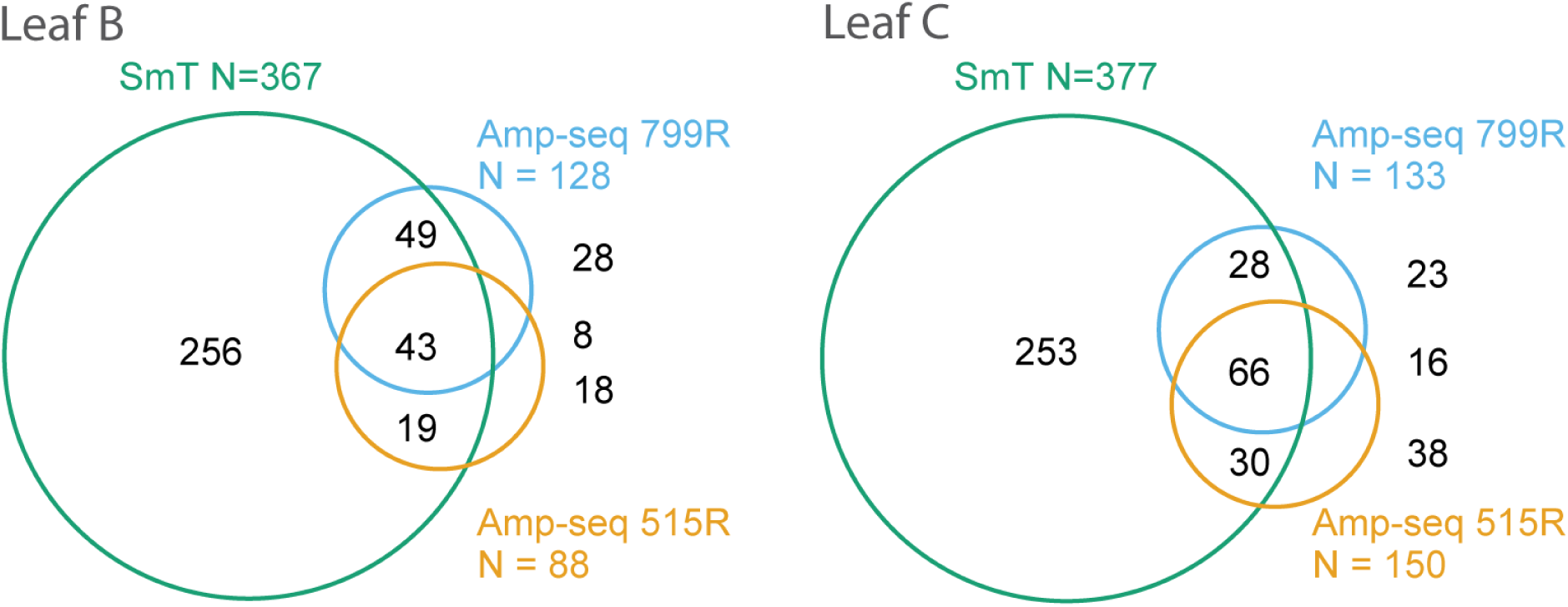
Taxa identified in different biological replicates presented in Venn diagrams. Values present the average of the subsetted samples after 100 iterations.

**Supplementary Figure S13:**
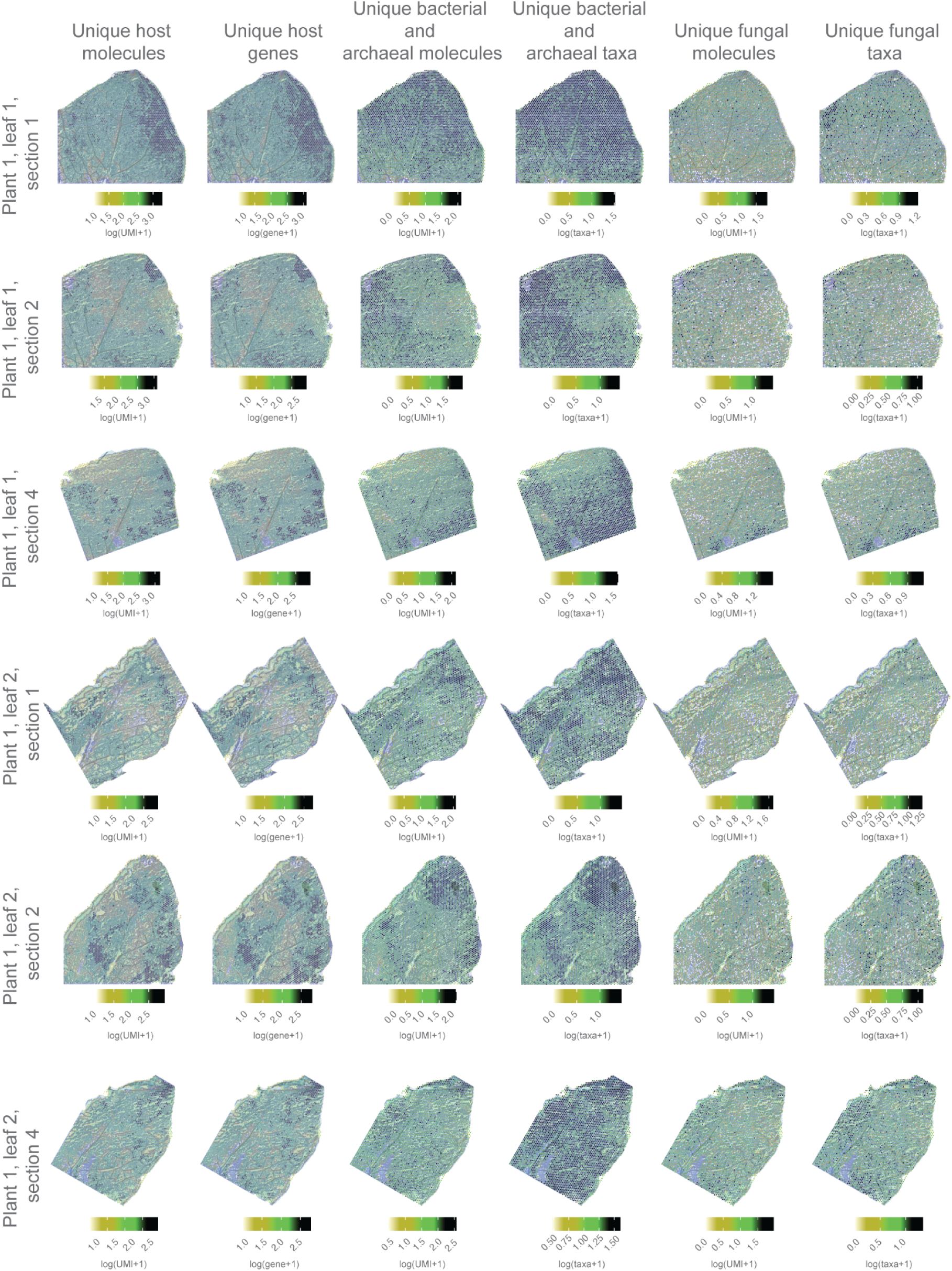
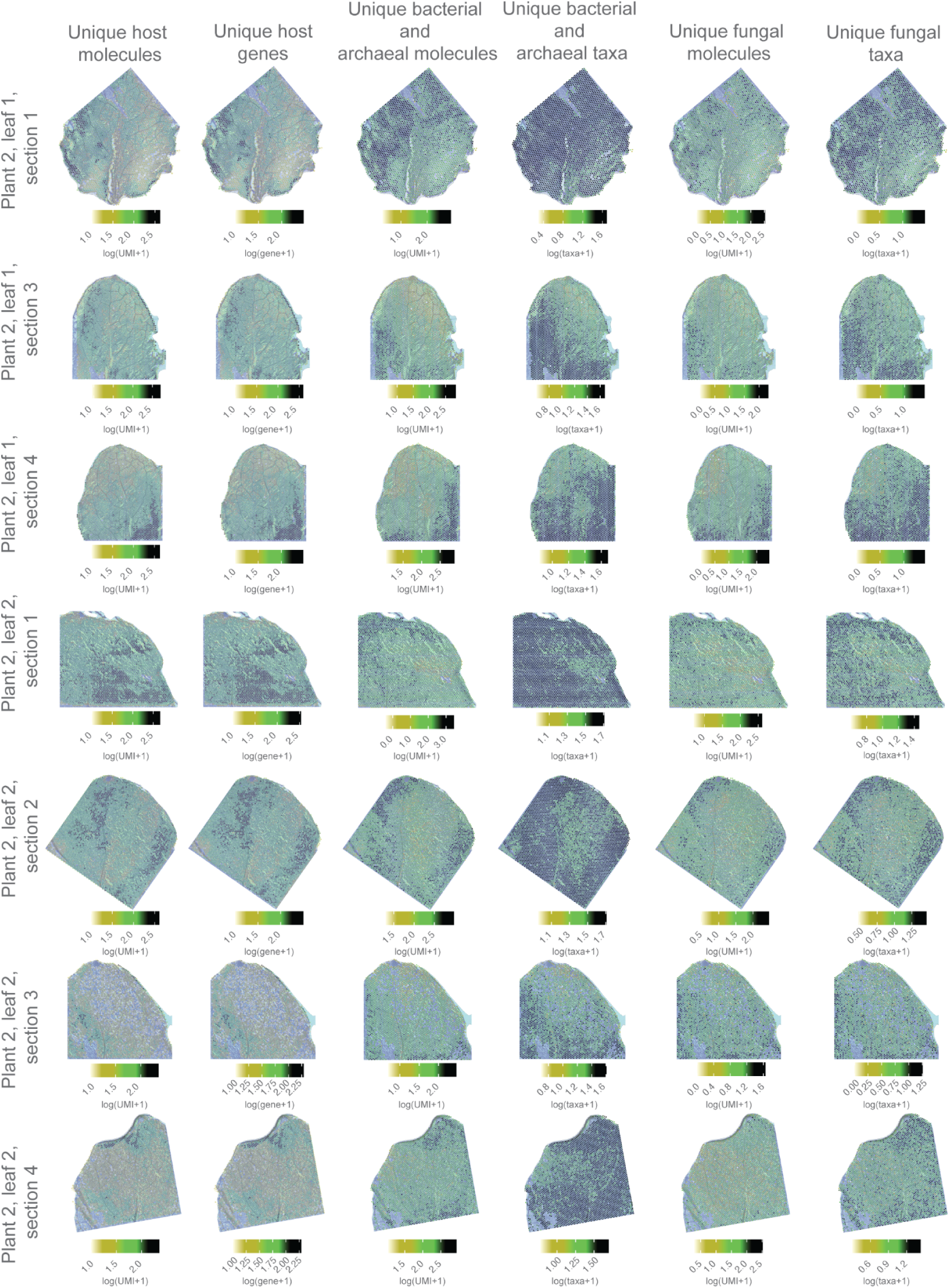
Unique host molecules, unique genes, unique bacterial and archaeal molecules, unique bacterial and archaeal taxa, unique fungal molecules and unique fungal taxa in log10 scale for all the sections in the dataset

**Supplementary Figure S14:**
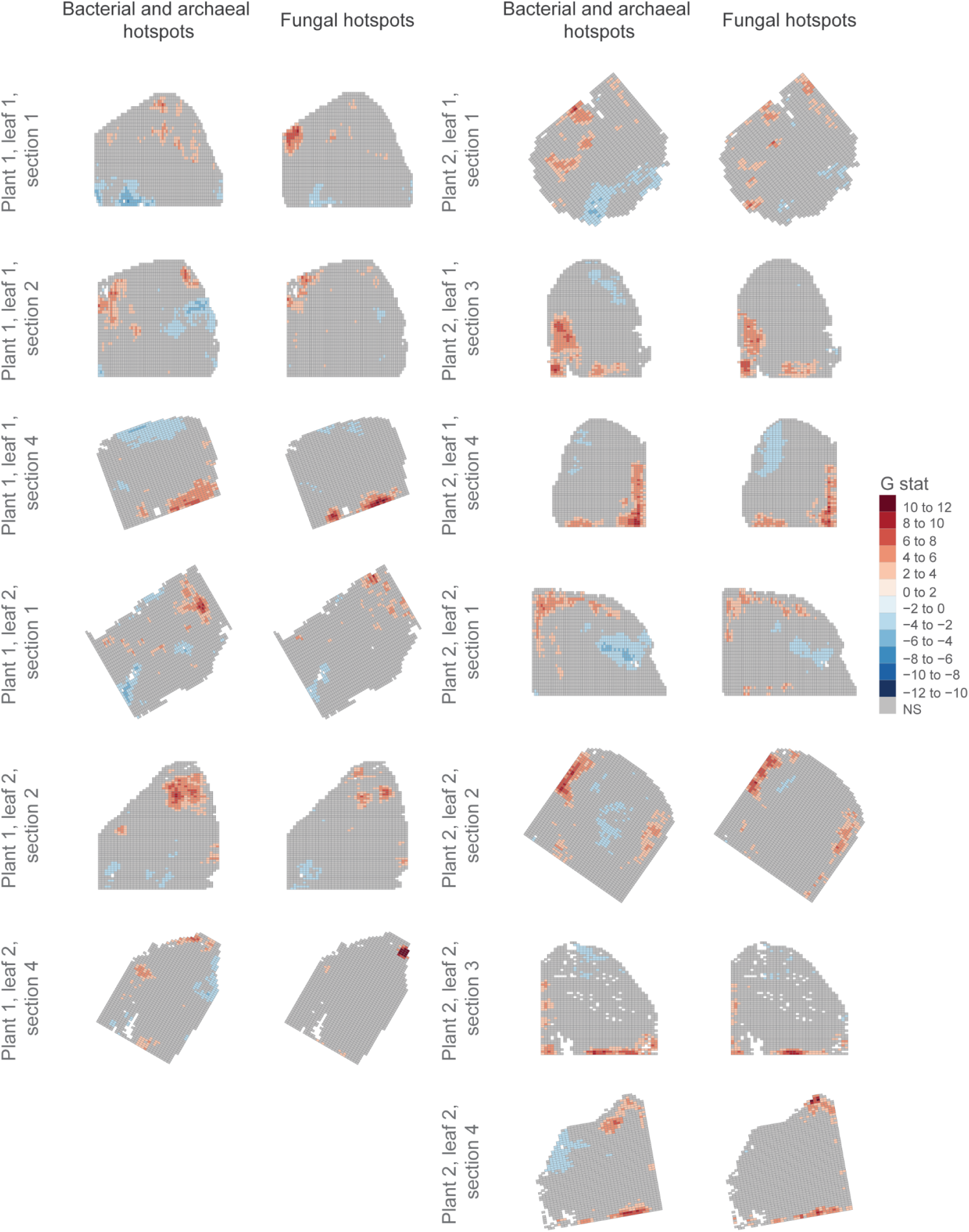
Bacterial and archaeal and fungal hotspots for each of the sections.

**Supplementary Figure S15:**
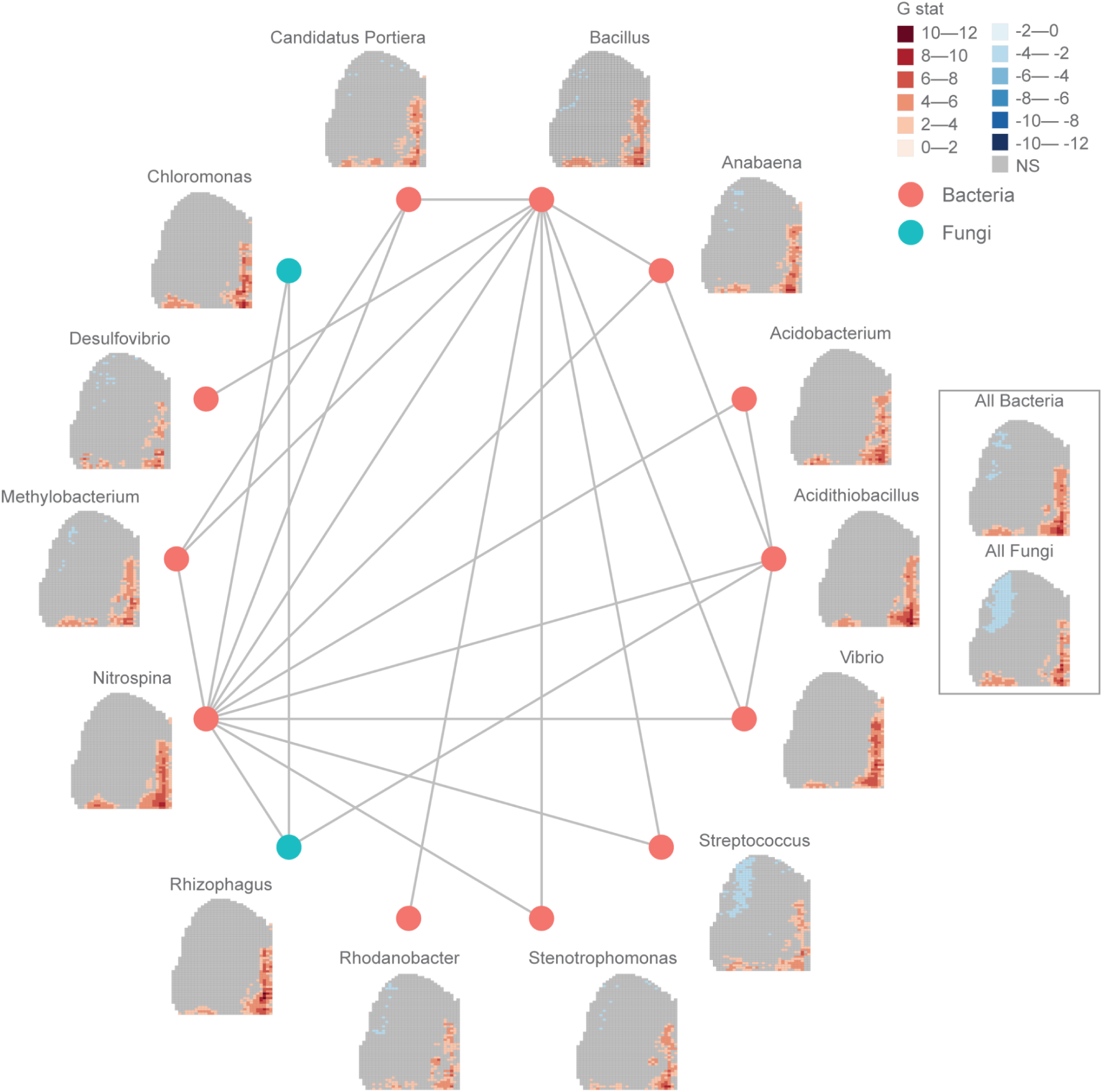
Subnetwork of 14 bacterial and fungal taxa strongly associated in all tested leaf sections. An edge connects two taxa if their average pairwise SRCC, across all leaf sections, is above or equal to 0.35. The hotspot pattern for each genus in a representative leaf section (P2.L1.4) is shown next to the network nodes.

**Supplementary Figure S16:**
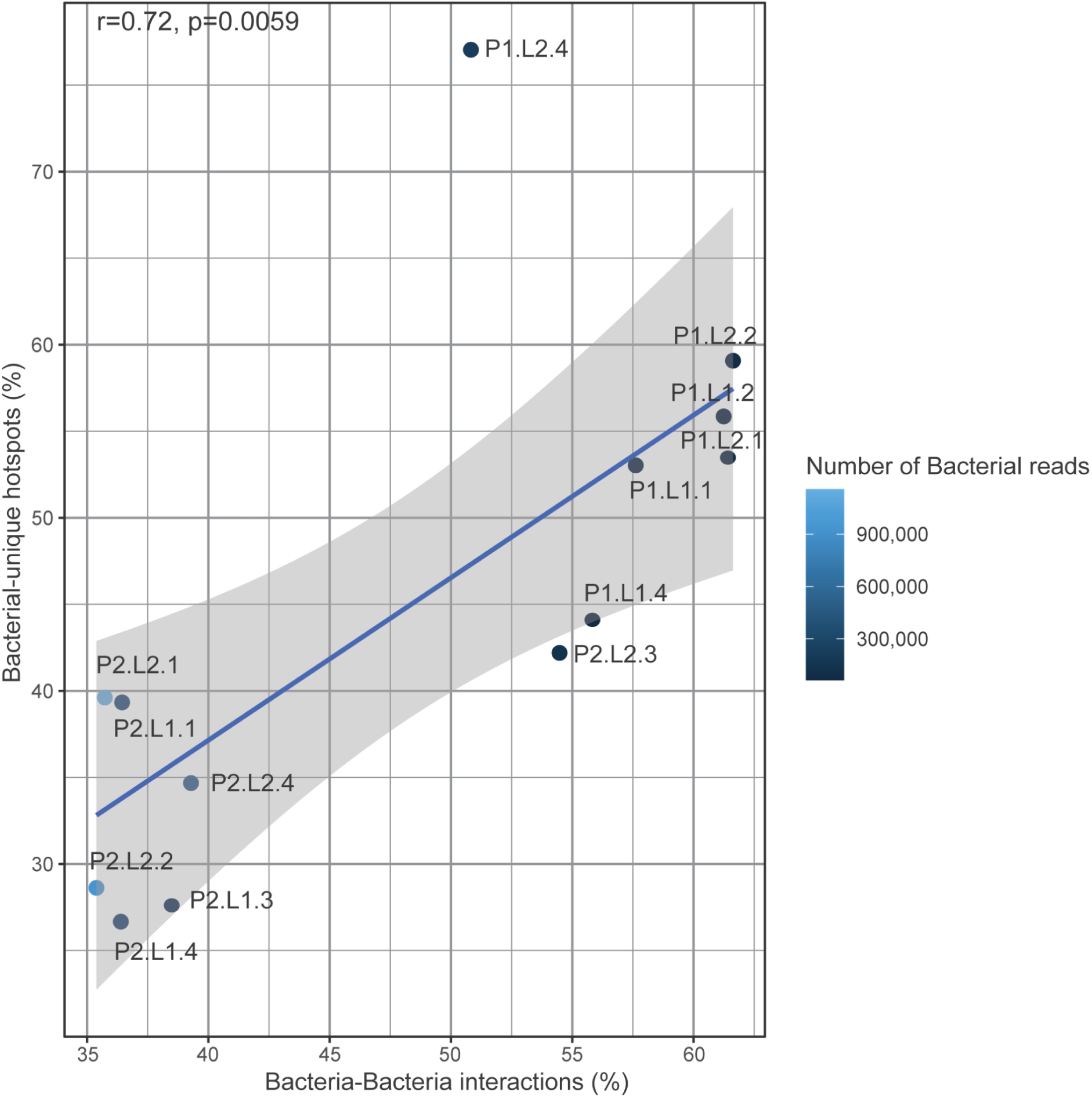
The proportion of bacteria-bacteria interactions as a function of the proportion of bacterial-unique hotspots.

**Supplementary Figure S17:**
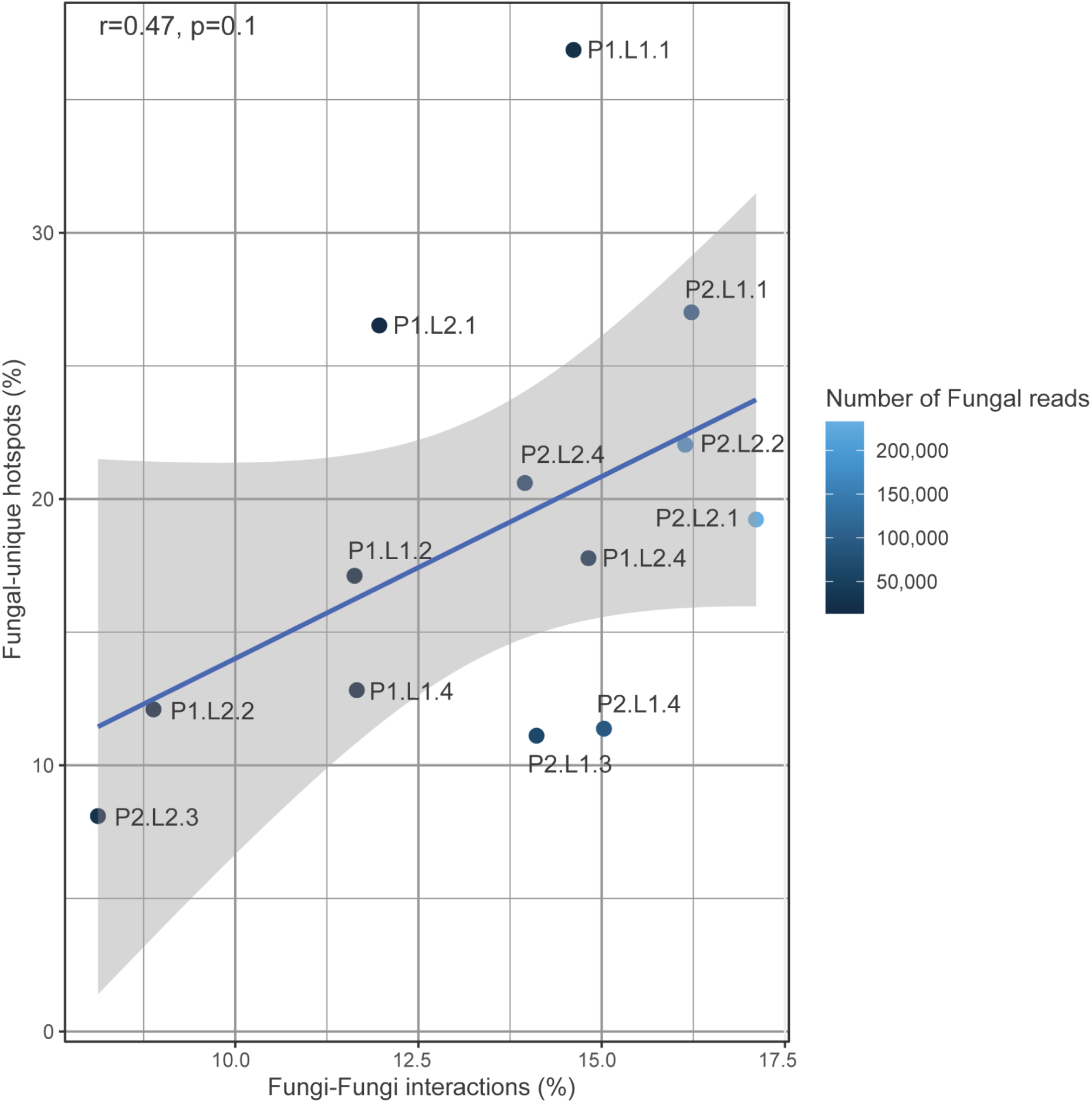
The proportion of fungi-fungi interactions as a function of the proportion of fungi-unique hotspots.

**Supplementary Figure S18:**
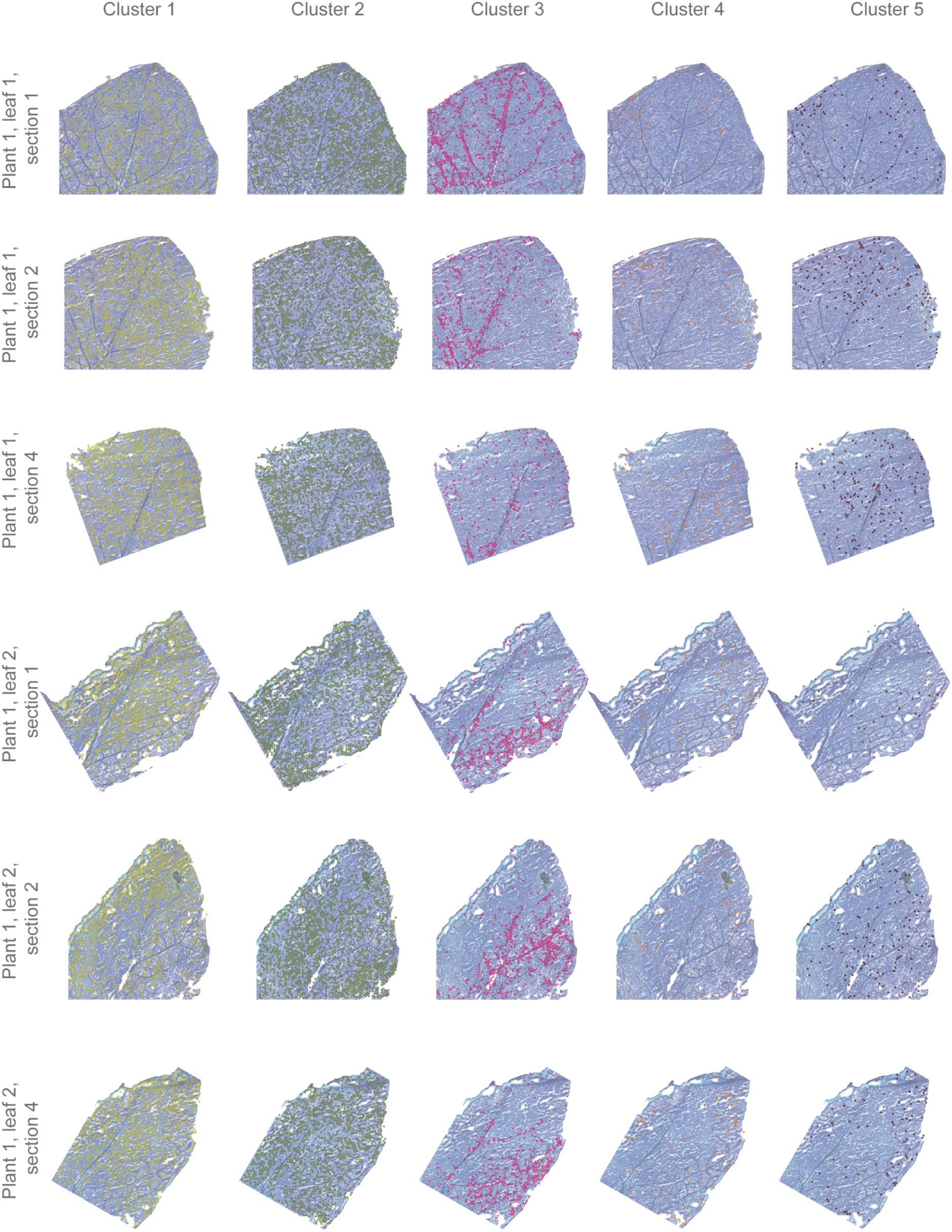
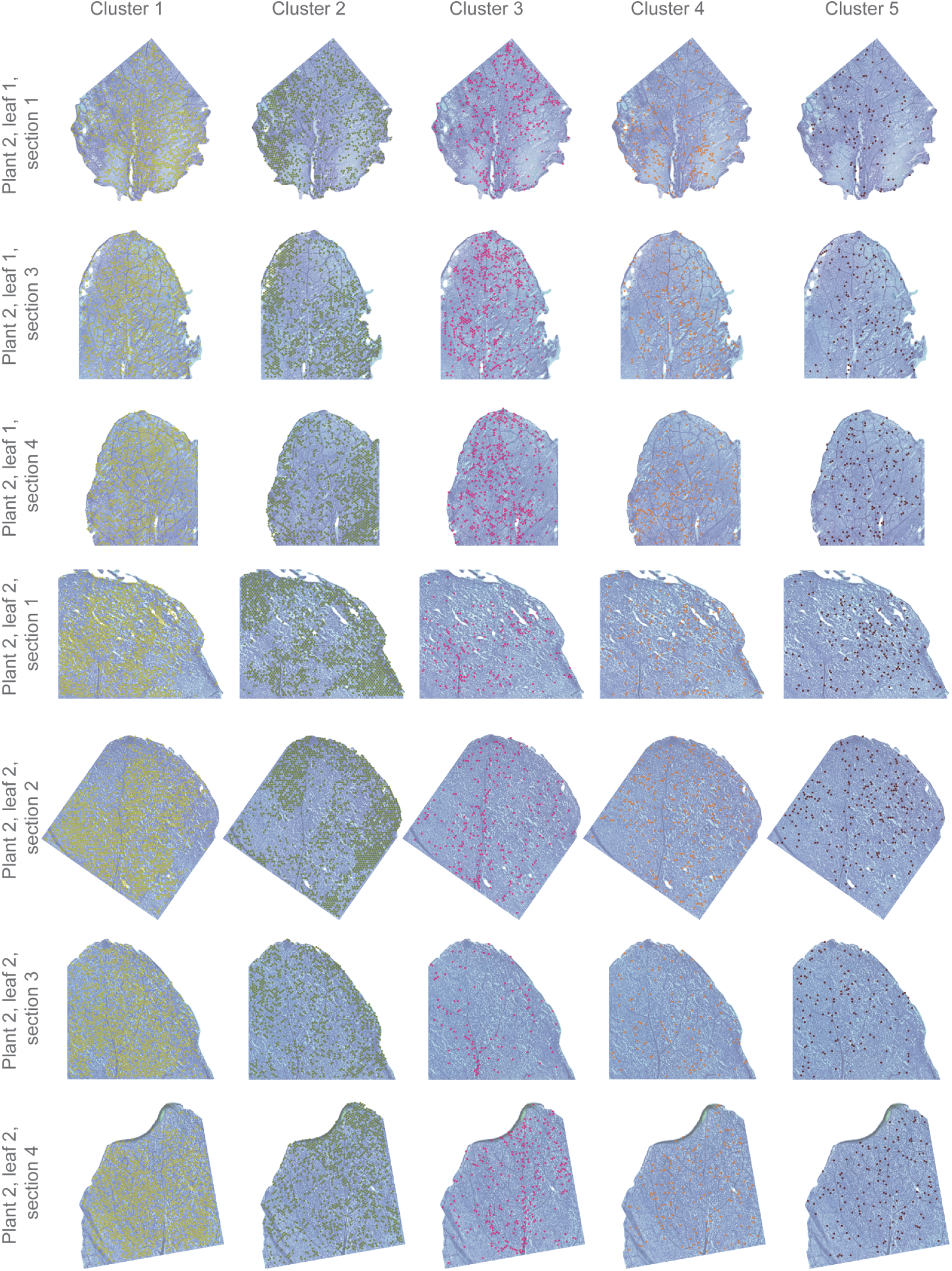
Each UMAP cluster is visualized separately in each tissue section.

**Supplementary Figure S19:**
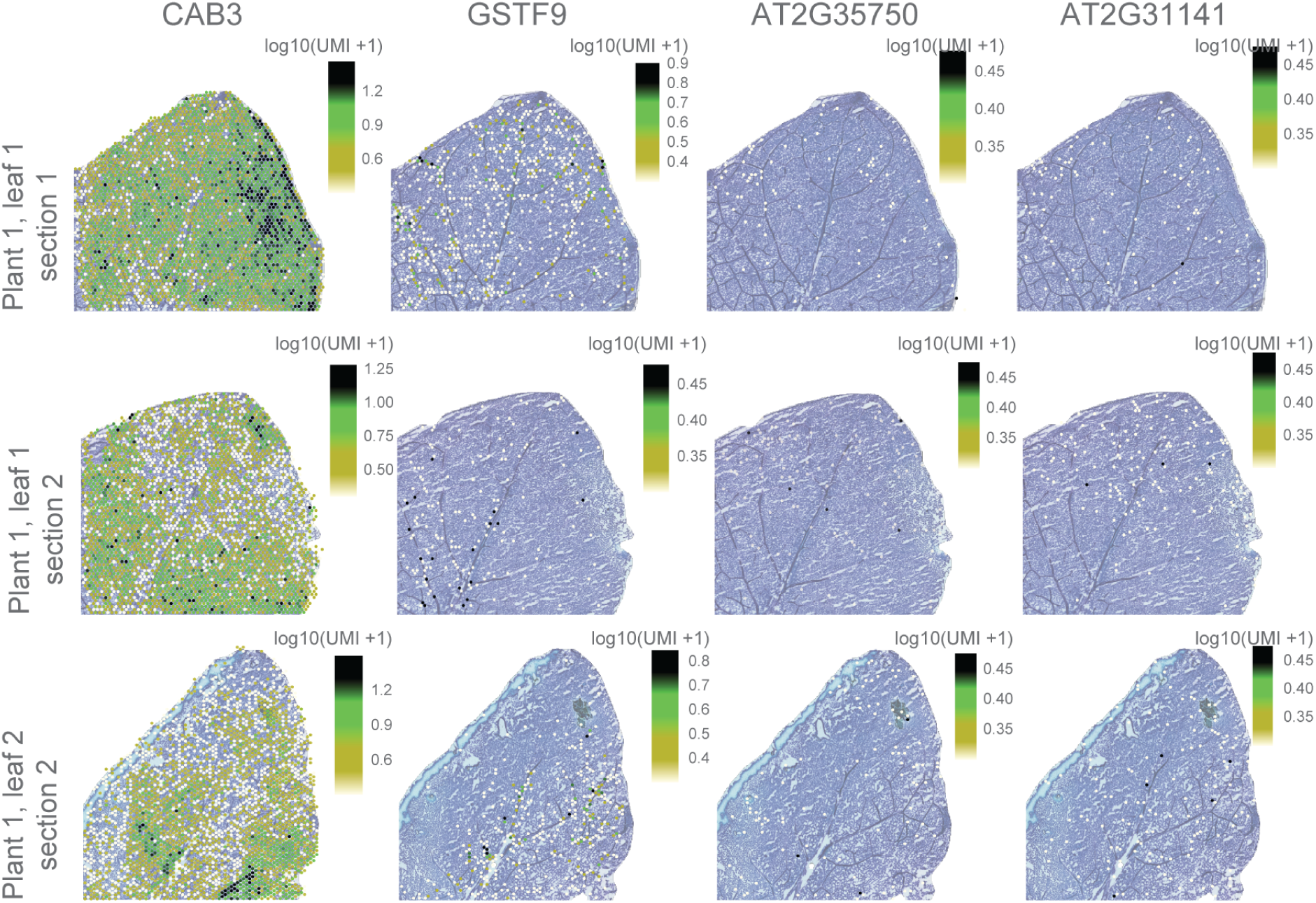
Representative markers for some of the clusters for three representative sections. CAB3: CHLOROPHYLL A/B BINDING PROTEIN 3 (Cluster 2 marker), GSTF9: GLUTATHIONE S-TRANSFERASE PHI 9 (Cluster 3 marker), AT2G35750: transmembrane protein (Cluster 4 marker), AT2G31141: potential nitrate responsive gene (Cluster 5 marker).

**Supplementary Figure S20:**
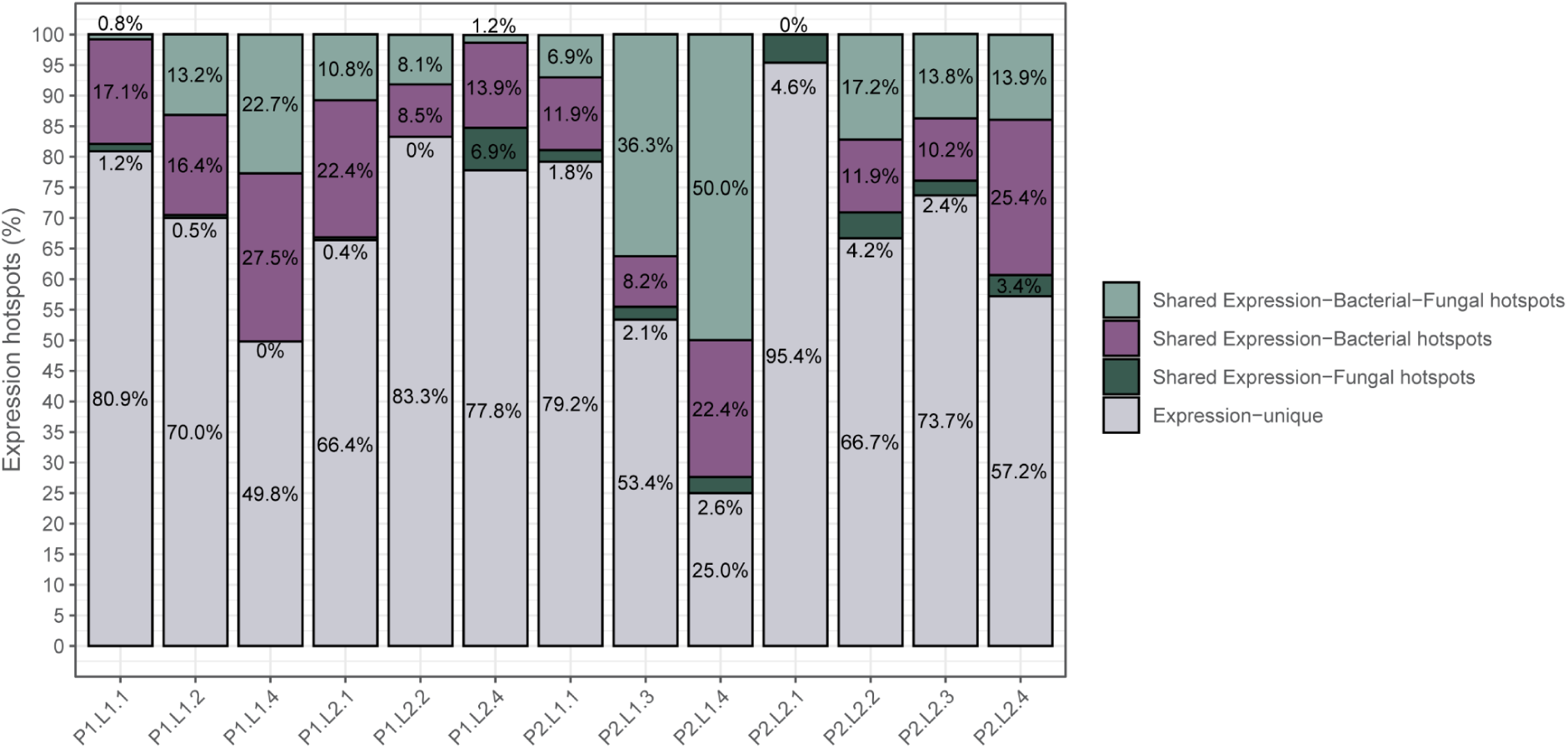
Proportion of hotspots shared between the host *A. thaliana*, bacterial and fungal organisms in each of the sections.

**Supplementary Figure S21:**
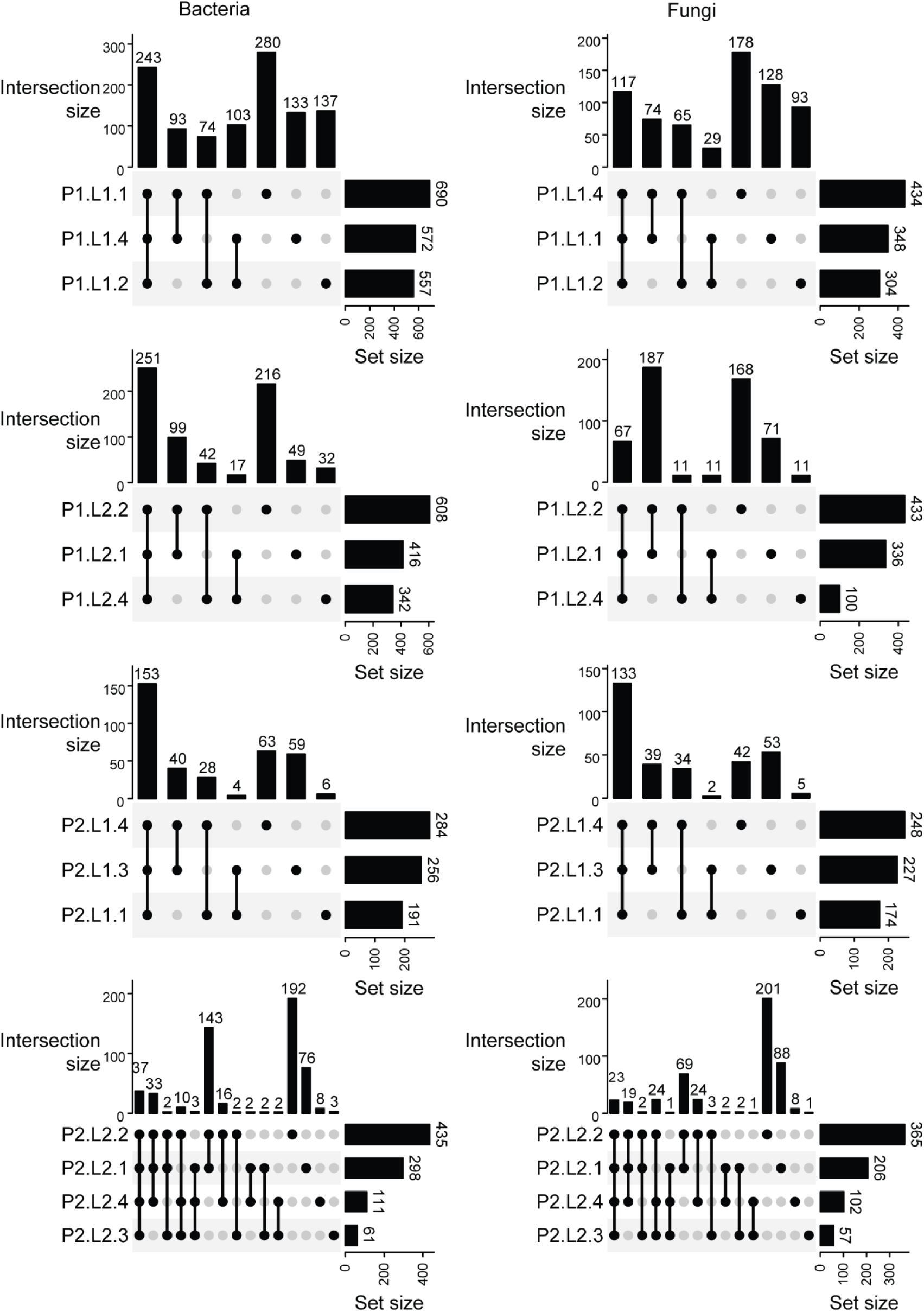
Set of genes selected by Boruta as explanatory of the total Bacterial or Fungi abundance in a given leaf section were intersected across the leaf sections. Intersection size of the different groups are shown.

**Supplementary Figure S22:**
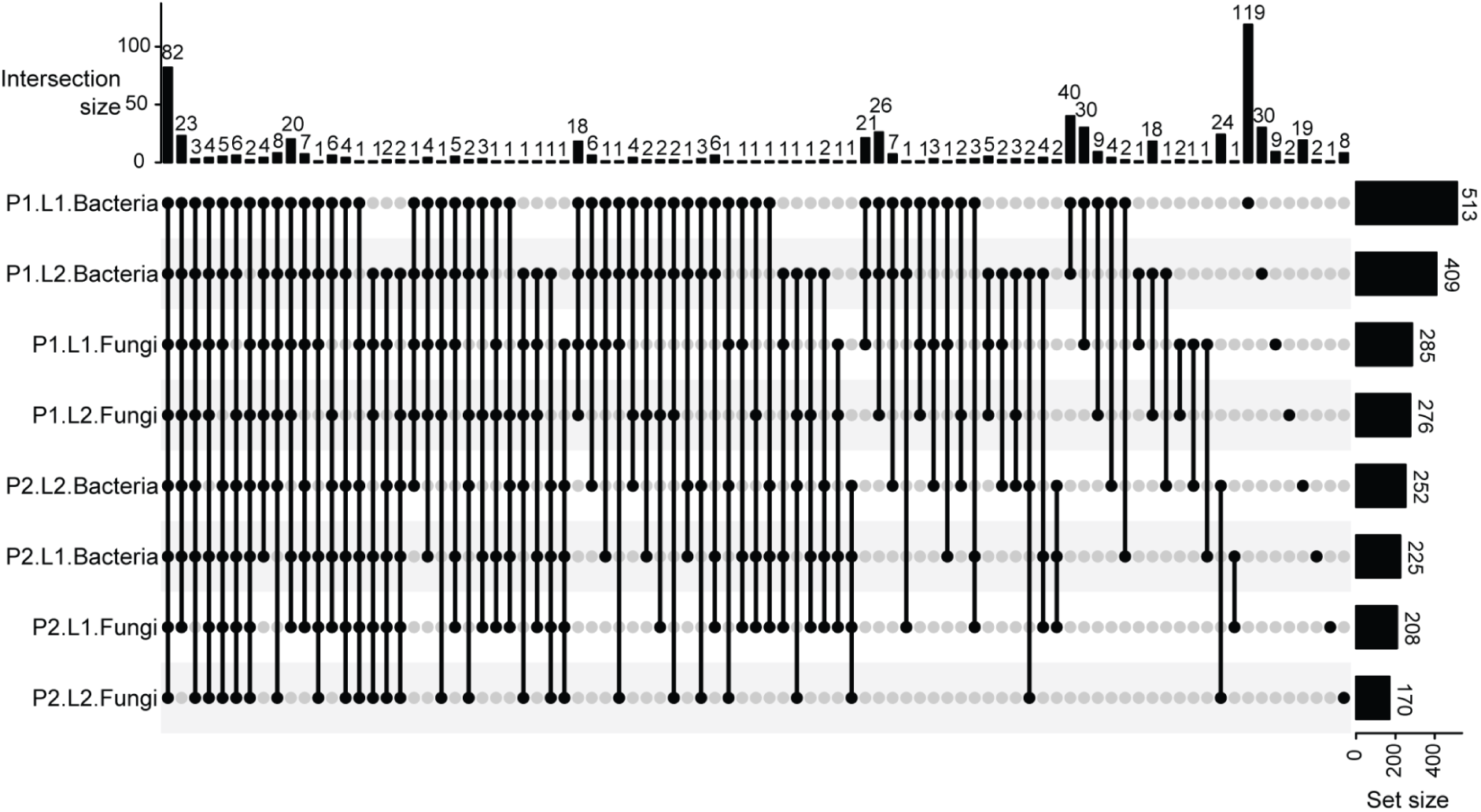
Intersection of Bacteria and Fungi associated genes across different leaves. Most of the genes are common to Bacteria and Fungi.

**Supplementary Figure S23:**
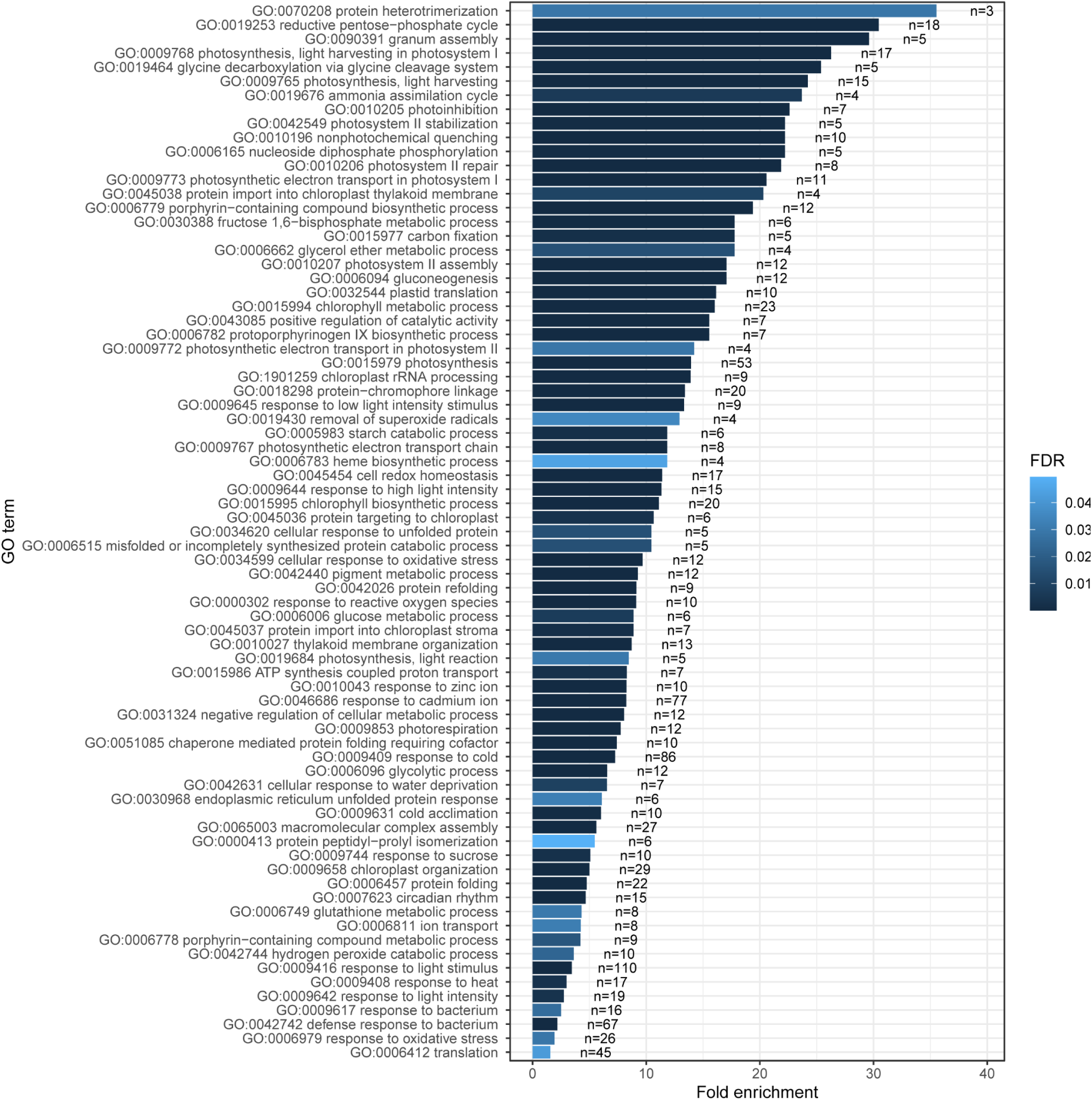
Overrepresented GO terms of all genes in the significantly associated with microbial abundance (n=the number of genes assigned with the specific go term).

**Supplementary Figure S24:**
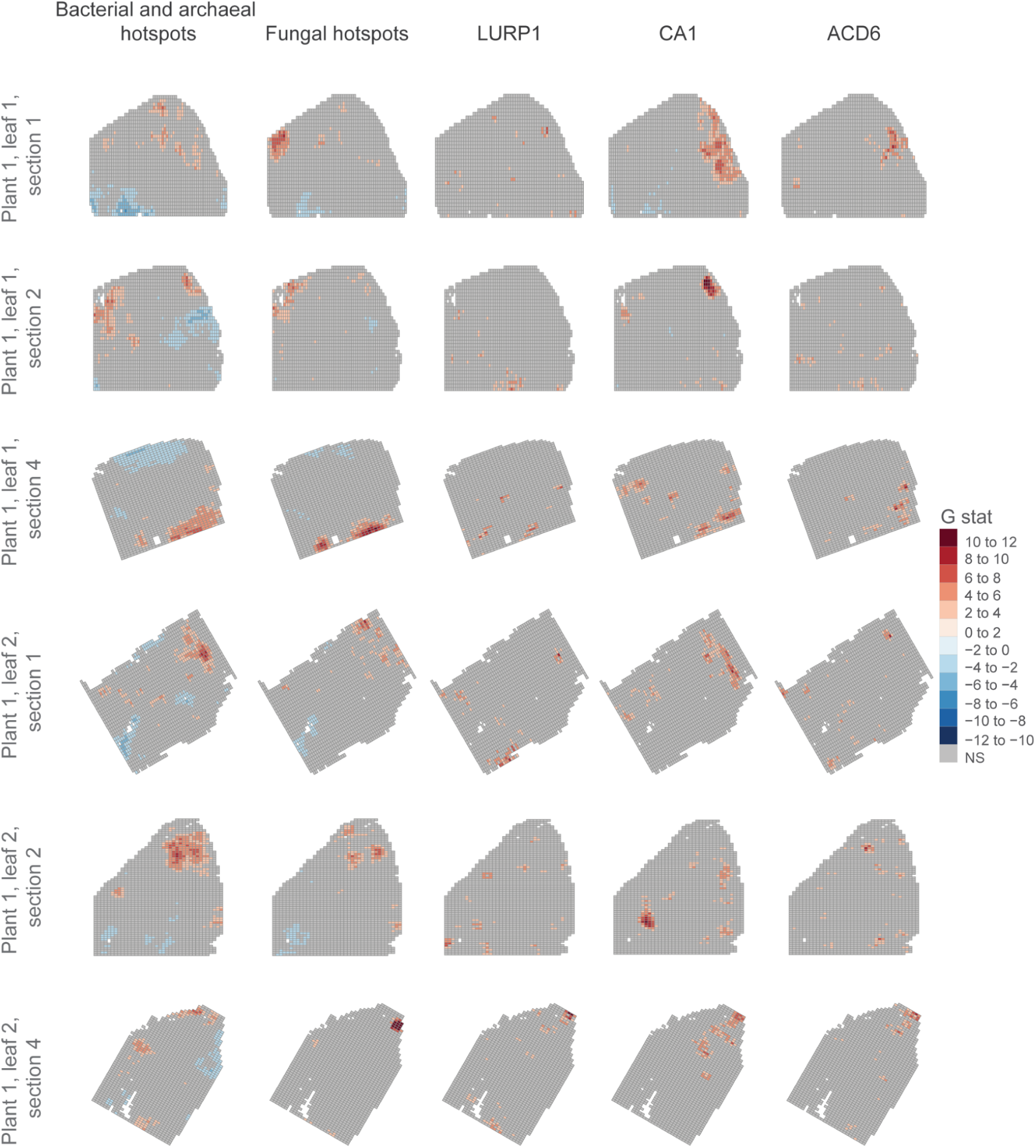
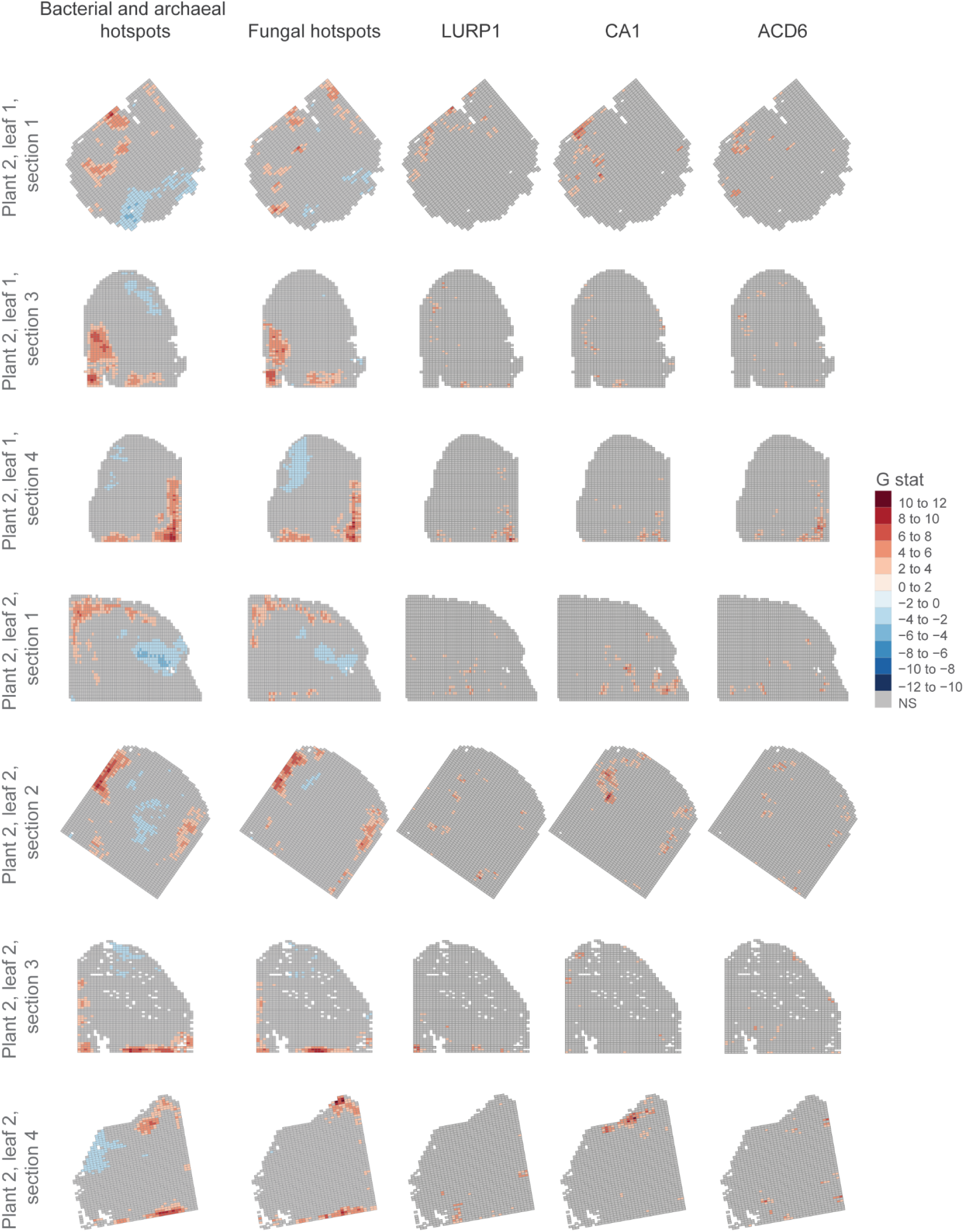
Spatial distribution of significant hotspots of bacteria, fungi, and three defense-related genes: *CA1* (AT3G01500), *LURP1* (AT2G14560), and *ACD6* (AT4G14400).

**Supplementary Table S1:** Bacterial, archeal and fungal taxa captured by different array types.

| **Array** | **Bacteria and archaea** |  |  | **Fungi** |  |  |
| --- | --- | --- | --- | --- | --- | --- |
|  | **Leaf 1** | **Leaf 2** | **Leaf 3** | **Leaf 1** | **Leaf 2** | **Leaf 3** |
| Multimodal array | 845 | 962 | 709 | 163 | 179 | 97 |
| 100% 16S array | 1,014 | 873 | 807 | 110 | 60 | 52 |
| 100% ITS/18S array | 494 | 566 | 356 | 304 | 304 | 253 |
| 100% poly-T array | 479 | 712 | 324 | 38 | 62 | 24 |

**Supplementary Table S2:** Bacterial and archeal relative abundance profile across samples.

| **Genus** | **Mean** | **SD** | **Median** |
| --- | --- | --- | --- |
| Nitrospina | 14.40% | 2.93% | 14.40% |
| Bacillus | 8.90% | 0.75% | 8.97% |
| Acidithiobacillus | 6.61% | 1.71% | 6.56% |
| Methylobacterium | 5.56% | 0.56% | 5.62% |
| Streptococcus | 4.69% | 0.46% | 4.70% |
| Streptomyces | 3.87% | 1.23% | 3.86% |
| Candidatus Portiera | 3.65% | 0.36% | 3.73% |
| Stenotrophomonas | 2.75% | 0.25% | 2.85% |
| Anabaena | 2.46% | 0.23% | 2.49% |
| Rhodanobacter | 2.36% | 0.22% | 2.38% |
| Vibrio | 2.12% | 0.35% | 2.17% |
| Desulfovibrio | 1.97% | 0.15% | 1.97% |
| Bradyrhizobium | 1.53% | 0.12% | 1.55% |
| Acinetobacter | 1.44% | 0.20% | 1.40% |
| Acidobacterium | 1.23% | 0.42% | 1.38% |
| Priestia | 1.19% | 0.15% | 1.19% |
| Mycoplasma | 1.14% | 0.30% | 1.03% |
| Pseudomonas | 1.02% | 0.10% | 1.03% |
| Clostridium | 1.01% | 0.31% | 0.90% |
| Marinobacter | 1.01% | 0.22% | 1.03% |
| Smithella | 0.99% | 0.37% | 0.84% |
| Geobacter | 0.85% | 0.15% | 0.82% |
| Halospirulina | 0.75% | 0.33% | 0.69% |
| Rhodobium | 0.73% | 0.51% | 0.53% |
| Lactobacillus | 0.68% | 0.21% | 0.67% |
| Cyanothece | 0.61% | 0.28% | 0.62% |
| Flavobacterium | 0.56% | 0.20% | 0.54% |
| Shewanella | 0.46% | 0.40% | 0.29% |
| Proteus | 0.40% | 0.32% | 0.25% |
| Other (<=1%) | 25.10% | 1.76% | 25.20% |

**Supplementary Table S3:** Fungal relative abundance profile across samples.

| **Genus** | **Mean** | **SD** | **Median** |
| --- | --- | --- | --- |
| Phaeosphaeria | 20.20% | 7.64% | 22.40% |
| Rhizophagus | 13.80% | 2.89% | 12.40% |
| Chloromonas | 11.70% | 2.06% | 11.60% |
| Amanita | 5.68% | 1.17% | 6.04% |
| Femsjonia | 5.51% | 1.63% | 5.22% |
| Elaphomyces | 4.62% | 0.99% | 4.78% |
| Roccella | 3.11% | 0.73% | 2.82% |
| Helminthosporium | 2.66% | 0.55% | 2.71% |
| Marasmius | 2.02% | 0.44% | 1.88% |
| Geopora | 1.92% | 0.45% | 1.86% |
| Elmerina | 1.71% | 0.36% | 1.61% |
| Plasmodium | 1.57% | 0.96% | 1.15% |
| Willeya | 1.49% | 0.38% | 1.44% |
| Phlyctochytrium | 1.04% | 0.19% | 1.06% |
| Aspergillus | 0.96% | 0.48% | 0.83% |
| Saccharomyces | 0.94% | 0.52% | 0.89% |
| Talaromyces | 0.88% | 0.17% | 0.84% |
| Strongyloides | 0.79% | 0.51% | 0.49% |
| Phellinus | 0.78% | 0.19% | 0.79% |
| Malassezia | 0.59% | 0.16% | 0.56% |
| Dictyochloris | 0.51% | 1.66% | 0.01% |
| Trichophyton | 0.44% | 0.22% | 0.39% |
| Phyllozyma | 0.44% | 0.37% | 0.33% |
| Other (<=1% in all samples) | 16.70% | 5.22% | 14.70% |

**Supplementary Table S4:** Array coverage statistics.

| Sample | Pixels<br>under the<br>tissue | Bacterial<br>pixels<br>under the<br>tissue | Fungi<br>pixels<br>under the<br>tissue | Bacteria<br>coverage | Fungi<br>coverage | Median<br>Bacterial<br>reads per<br>pixel<br>(RpM) | Median<br>Fungi<br>reads per<br>pixels<br>(RpM) | Median<br>mRNA<br>reads per<br>pixel<br>(RpM) | Median<br>Bacterial<br>reads per<br>pixels | Median<br>Fungal<br>reads per<br>pixel | Median<br>mRNA<br>reads per<br>pixel |
| --- | --- | --- | --- | --- | --- | --- | --- | --- | --- | --- | --- |
| P1.L1.1 | 3,451 | 3,444 | 3,330 | 99.8% | 96.5% | 276.759 | 227.097 | 248.997 | 32.0 | 5.0 | 314.0 |
| P1.L1.2 | 3,730 | 3,727 | 3,403 | 99.9% | 91.2% | 252.323 | 225.513 | 215.385 | 17.0 | 3.0 | 136.5 |
| P1.L1.4 | 2,997 | 2,990 | 2,815 | 99.8% | 93.9% | 298.105 | 275.406 | 216.516 | 19.0 | 4.0 | 103.0 |
| P1.L2.1 | 3,561 | 3,555 | 3,313 | 99.8% | 93.0% | 254.155 | 256.674 | 208.034 | 20.0 | 4.0 | 72.0 |
| P1.L2.2 | 3,704 | 3,700 | 3,469 | 99.9% | 93.7% | 234.098 | 242.557 | 172.493 | 18.0 | 4.0 | 91.0 |
| P1.L2.4 | 3,074 | 3,074 | 3,064 | 100.0% | 99.7% | 310.658 | 250.708 | 274.790 | 66.0 | 10.0 | 103.5 |
| P2.L1.1 | 2,765 | 2,765 | 2,763 | 100.0% | 99.9% | 332.178 | 298.530 | 292.632 | 143.0 | 31.0 | 53.0 |
| P2.L1.3 | 2,897 | 2,897 | 2,896 | 100.0% | 100.0% | 257.571 | 276.919 | 293.053 | 64.0 | 18.0 | 77.0 |
| P2.L1.4 | 2,725 | 2,725 | 2,721 | 100.0% | 99.9% | 287.726 | 302.768 | 294.028 | 108.0 | 29.0 | 63.0 |
| P2.L2.1 | 3,485 | 3,485 | 3,485 | 100.0% | 100.0% | 265.394 | 253.449 | 258.088 | 307.0 | 59.0 | 117.0 |
| P2.L2.2 | 4,083 | 4,083 | 4,083 | 100.0% | 100.0% | 218.112 | 201.729 | 209.358 | 206.0 | 36.0 | 80.0 |
| P2.L2.3 | 2,731 | 2,731 | 2,718 | 100.0% | 99.5% | 331.450 | 320.936 | 269.929 | 46.0 | 9.0 | 19.0 |
| P2.L2.4 | 3,186 | 3,186 | 3,186 | 100.0% | 100.0% | 284.079 | 265.382 | 255.544 | 153.0 | 20.0 | 28.0 |

**Supplementary Table S5:**
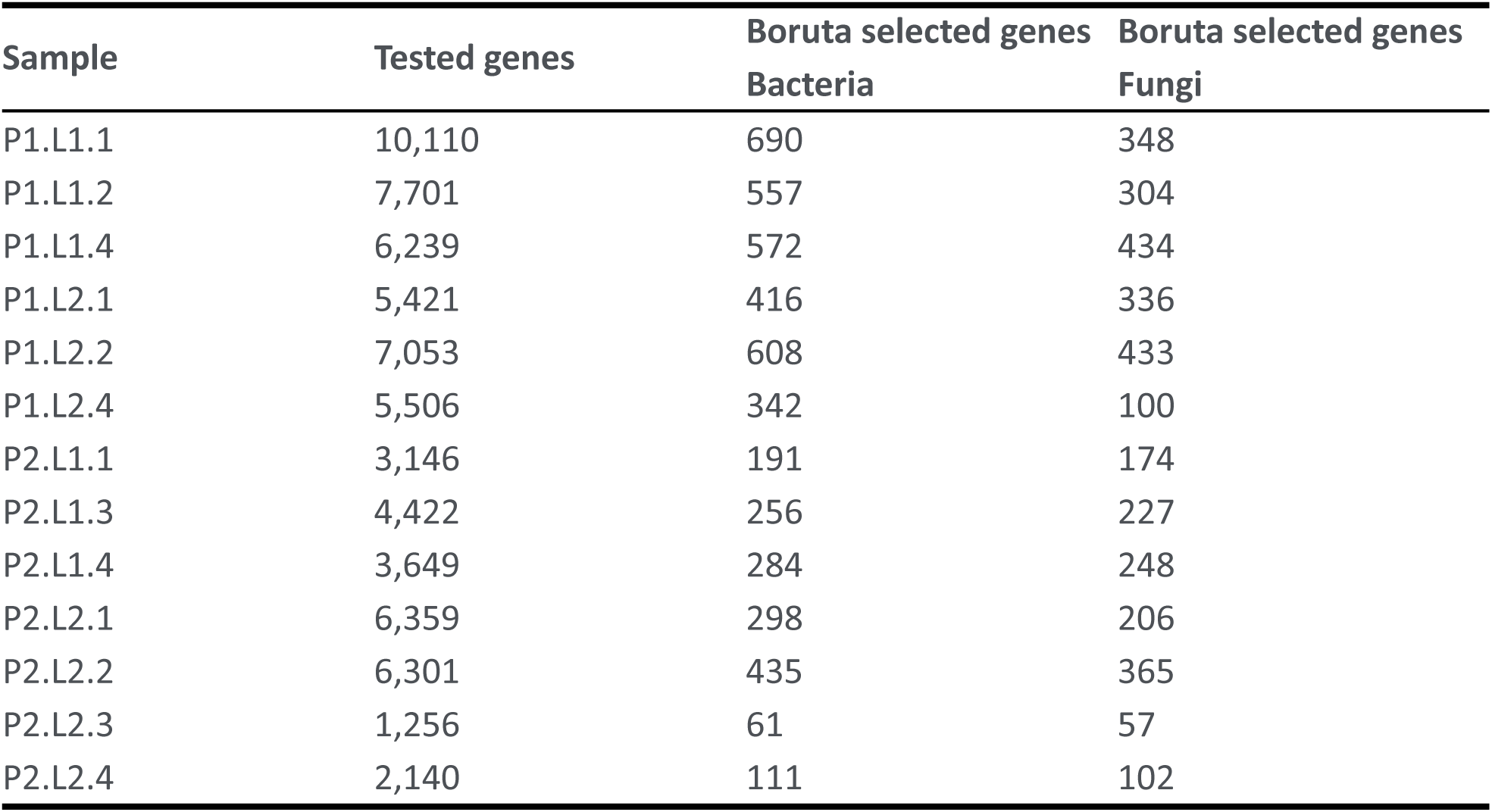
Number of genes selected by Boruta.

**Supplementary Table S8:** Number of annotated reads considered for each dataset.

| Sample | Bacterial reads | Bacterial read (top 50 genera) | Bacterial reads under the tissue (top 50 genera) | Fungi reads | Fungi read (top 50 genera) | Fungi reads under the tissue (top 50 genera) | mRNA reads |
| --- | --- | --- | --- | --- | --- | --- | --- |
| P1.L1.1 | 181,887 | 153,548 | 115,624 | 36,992 | 32,915 | 22,017 | 1,261,057 |
| P1.L1.2 | 104,104 | 86,223 | 67,374 | 22,050 | 18,296 | 13,303 | 633,750 |
| P1.L1.4 | 102,844 | 85,738 | 63,736 | 24,411 | 20,748 | 14,524 | 475,715 |
| P1.L2.1 | 117,185 | 96,829 | 78,692 | 25,379 | 21,059 | 15,584 | 346,097 |
| P1.L2.2 | 117,244 | 95,783 | 76,891 | 26,123 | 21,861 | 16,491 | 527,559 |
| P1.L2.4 | 329,956 | 280,809 | 212,452 | 69,162 | 61,984 | 39,887 | 376,651 |
| P2.L1.1 | 641,002 | 545,808 | 430,492 | 139,230 | 129,020 | 103,842 | 181,115 |
| P2.L1.3 | 364,618 | 301,947 | 248,475 | 96,852 | 86,505 | 65,001 | 262,751 |
| P2.L1.4 | 550,994 | 459,428 | 375,357 | 127,954 | 115,187 | 95,783 | 214,265 |
| P2.L2.1 | 1,611,258 | 1,388,598 | 1,156,771 | 289,374 | 274,587 | 232,788 | 453,334 |
| P2.L2.2 | 1,243,355 | 1,060,542 | 944,468 | 213,149 | 199,767 | 178,457 | 382,121 |
| P2.L2.3 | 219,430 | 177,920 | 138,784 | 44,088 | 37,470 | 28,043 | 70,389 |
| P2.L2.4 | 824,179 | 691,206 | 538,583 | 115,987 | 101,169 | 75,363 | 109,570 |

